# Excess Met1-ubiquitination leads to solid aggregate formation

**DOI:** 10.64898/2026.01.20.700516

**Authors:** Stephanie Kaypee, Minori Miyasaka, Takuto Nakajima, Hiromi Nishimura, Jun-ichi Sakamaki, Masaaki Komatsu, Fumiyo Ikeda

## Abstract

The ubiquitin ligase HOIL-1 has a unique role in controlling quantity of Met1-linked/ linear ubiquitin chains in cells by coordinating action with the ubiquitin ligase HOIP. Both ligases are components of the Linear UBiquitin chain Assembly Complex (LUBAC), the only known ligase complex that is able to generate Met1-linked ubiquitin chains. Although importance of Met1-linked ubiquitin chains in inflammation and immunity is well established, physiological relevance of quantity of these chains remain unknown. Here, we demonstrate that cells expressing catalytically inactive HOIL-1 exhibited significantly higher numbers of α-Synuclein, tau, and amyloid beta aggregates. This phenotype is associated with a disruption in late-stage autophagic flux, wherein p62-positive aggregates fail to colocalize with lysosomal markers, leading to impaired clearance. Additionally, a biophysical transition in aggregate properties was observed *in vitro*, with mutant cells forming more rigid solid-like inclusions, shifting from dynamic and liquid-like condensates. Elevated Met1-linked ubiquitin chains, either through HOIL-1 catalytic inactivation or knockdown of the Met1-linked chain-specific deubiquitinase OTULIN, phenocopied the defects in aggregate clearance. These findings reveal a critical role of HOIL-1 catalytic activity in modulating aggregate clearance through autophagy and maintaining the quantity of Met1-ubiquitin chains, highlighting HOIL-1 as a key factor in proteostasis in neurodegenerative diseases.

**Highlights:**

- HOIL-1 catalytic activity prevents neurodegenerative protein aggregate accumulation
- Inactive HOIL-1 impairs late-stage autophagic clearance of protein aggregates
- Loss of HOIL-1 shifts aggregates to rigid, solid-like states
- Excess Met1-ubiquitin chains drive aggregate solidification

## Introduction

Proteostasis, the maintenance of protein homeostasis, is crucial for ensuring proper protein synthesis, folding, trafficking and degradation (Bauer *et al*, 2023; Sala *et al*, 2017). Failure of proteostasis results in protein misfolding and aggregation (Ciechanover & Kwon, 2015; Sala *et al*., 2017). Protein aggregation is a characteristic feature of neurodegenerative disorders such as Alzheimer’s disease (AD), Parkinson’s disease (PD), and amyotrophic lateral sclerosis (ALS) (Moda *et al*, 2023). Accumulation of misfolded proteins in neurodegenerative diseases, including α-Synuclein, tau, amyloid-β (Aβ), and TAR DNA-binding protein 43 (TDP-43), disrupts cellular homeostasis and leads to neuronal dysfunction (Ciechanover & Kwon, 2015; Moda *et al*., 2023). Cells employ specialized pathways to manage protein aggregates, such as degradation via the ubiquitin-proteasome system (UPS) and autophagic clearance (Kocaturk & Gozuacik, 2018). The UPS targets misfolded or damaged ubiquitin-tagged proteins for degradation through the 26S proteasome (Hipp *et al*, 2019); however, its ability to degrade large aggregates is limited (Kocaturk & Gozuacik, 2018). Autophagy complements the UPS by engulfing protein aggregates into autophagosomes, which fuse with lysosomes for degradation (Dikic, 2017).

While ubiquitination is a key signal for both the UPS and autophagy, the specific types and topologies of ubiquitin chains play distinct roles in these processes (Agrata & Komander, 2025; Akizuki *et al*, 2024). Of particular interest in proteostasis and aggregate clearance are Met1-linked ubiquitin chains, whose critical yet underexplored involvement in cellular quality control and neurodegenerative contexts has recently been highlighted (Bellenguez *et al*, 2022; Furthmann *et al*, 2023; Tokunaga & Ikeda, 2022; van Well *et al*, 2019). For example, a recent genetic study on the etiology of AD and related dementias identified several key genes, including “Heme-oxidized IRP2 ubiquitin ligase 1 (HOIL-1)”, “Shank-associated RH domain-interacting protein (SHARPIN)”, “OTU domain-containing deubiquitinase with linear linkage specificity (OTULIN)”, and “TNFAIP3-interacting protein 1 (TNIP1)” (Bellenguez *et al*., 2022), all of which play roles in the Met1-linked ubiquitination system (Rittinger & Ikeda, 2017; Wagner *et al*, 2008). Met1-linked ubiquitin chains have been implicated in distinct signalling pathways, including inflammation(Fiil & Gyrd-Hansen, 2021; Jahan *et al*, 2021) and more recently, cellular quality control (Tokunaga & Ikeda, 2022), although their precise roles in modulating aggregate dynamics and clearance remain poorly understood. A comprehensive understanding of how Met1-linked ubiquitination, particularly through intricate regulation by Linear Ubiquitin Chain Assembly Complex (LUBAC) components, such as HOIL-1, influences aggregate dynamics and clearance; therefore, it is crucial to develop targeted therapeutic strategies against neurodegenerative proteinopathies.

LUBAC is responsible for forming these critical Met1-linked ubiquitin chains (Kirisako *et al*, 2006). LUBAC comprises the E3 ligase “HOIL-1-interacting protein (HOIP)”, which has a catalytic center for Met1 ubiquitination, another E3 ligase, HOIL-1, and the regulatory subunit SHARPIN (Gerlach *et al*, 2011; Ikeda *et al*, 2011; Tokunaga *et al*, 2011). HOIL-1 plays a unique regulatory role in this complex, which is crucial for Met1-linked ubiquitination (Kelsall *et al*, 2019). Its catalytic activity, which is involved in the formation of specific heterotypic ubiquitin chains, is essential for fine-tuning the overall levels of Met1-linked ubiquitin chains (Rodriguez Carvajal *et al*, 2021). Indeed, its catalytic inactivation leads to an increase in these chains, suggesting a suppressive or balancing function of HOIL-1 on Met1-linked ubiquitination activity of HOIP. A growing body of evidence highlights the diverse functions of LUBAC, extending from immune signal regulation to cellular quality control mechanisms, particularly the regulation of protein aggregation (Tokunaga & Ikeda, 2022). Recent findings suggest that Met1-linked ubiquitination is involved in marking misfolded or aggregated proteins and modulating stress responses (Asaoka *et al*, 2016; van Well *et al*., 2019). LUBAC components have been observed to localize to protein aggregates, and their activity appears to influence the clearance of aggregates through autophagy or proteasomal degradation (Furthmann *et al*., 2023; van Well *et al*., 2019). However, despite these observations, the specific contribution of HOIL-1 catalytic activity to these processes, particularly its impact on the material properties of aggregates and efficiency of autophagic flux, remains poorly defined. Specifically, three critical questions persist: (1) How does HOIL-1 catalytic activity specifically regulate aggregate clearance? (2) What is the relationship between the Met1-linked chain levels and the biophysical properties of aggregates? (3) How does this regulation affect autophagic flux?

To address these critical questions, we hypothesized that the catalytic activity of HOIL-1 within LUBAC is crucial for fine-tuning Met1-linked ubiquitination and that dysregulation of these chains directly affects the biophysical properties of aggregates, thereby preventing the accumulation of toxic protein aggregates and maintaining their dynamic liquid-like state to facilitate efficient autophagic clearance. To test this hypothesis, we systematically investigated the role of HOIL-1 catalytic activity in aggregate management using cellular models of neurodegeneration and *in vitro* biochemical reconstitution, with a focus on its impact on aggregate biophysical properties and autophagic flux.

## Results

### Aggregate-prone proteins accumulate amyloid in neuroblastoma cells expressing catalytically inactive HOIL-1 mutant

To determine whether HOIL-1 catalytic activity influences protein aggregation, we used SH-SY5Y (human neuroblastoma) cells, a widely recognized cellular model for studying neurodegenerative protein aggregation. We examined three well-established aggregation-prone proteins in these cells, α-Synuclein, tau, and Aβ. SH-SY5Y cells expressing the catalytically inactive HOIL-1 C460A mutant (where the ubiquitin-loading Cys is mutated to Ala) showed a significantly higher number of amyloid-positive aggregates for ⍺-Synuclein, tau, and Aβ compared to cells expressing wild-type HOIL-1 (Figure 1A-C). Notably, HOIL-1 C460A was detected within these structures, as demonstrated by its colocalization with tau aggregates (Figure 1D).

**Figure 1.**
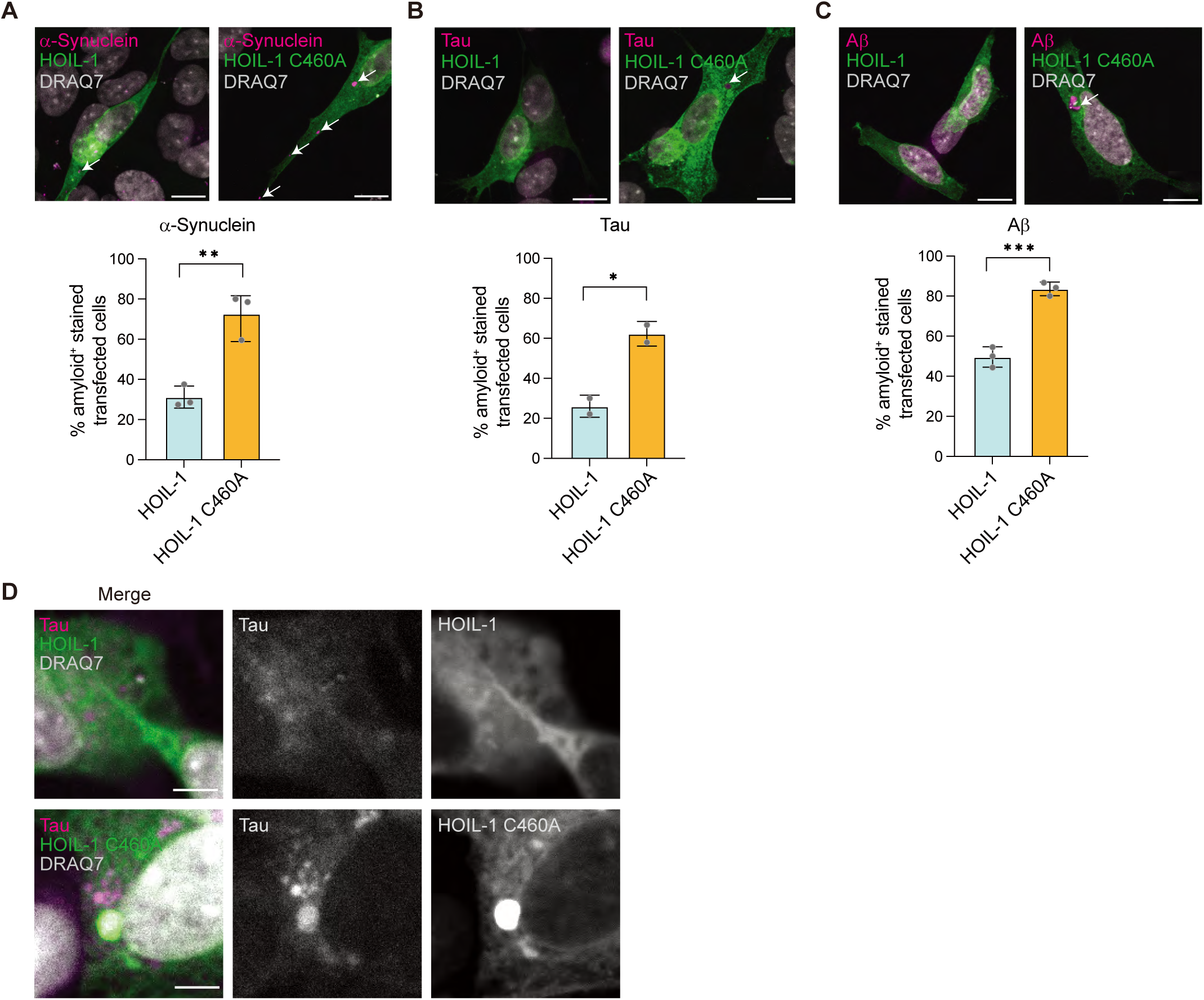
Aggregate-prone proteins accumulate in cells expressing catalytically inactive HOIL-1 C460A mutant. A-C. Comparison of α-Synuclein (A), tau (B), and Aβ (C) aggregates (indicated by arrows) in SH-SY5Y cells expressing GFP-HOIL-1 or GFP-HOIL-1 C460A. Scale bar = 10 μm. Graphical representations of the percentage of amyloid^+^ stained transfected cells are shown below. Mean ± SD was plotted, and unpaired t-test statistical analysis was performed; (A) **, P=0.0048, (B) *, P=0.0252, (C) ***, P=0.0006. (A) *n* ≥ 100 cells, N=3, (B) *n* ≥ 100 cells, N=2, (C) *n* ≥ 25 cells, N=3 independent biological repeats. D. Representative microscopy images showing colocalization of tau aggregates and GFP-tagged HOIL-1 or HOIL-1 C460A in SH-SY5Y cells. Scale bar = 5 μm. N=2 independent biological repeats.

To assess whether LUBAC components are associated with insoluble aggregates, we performed sucrose-gradient fractionation to separate insoluble aggregates from soluble protein fractions in SH-SY5Y cells co-expressing α-Synuclein with either HOIL-1 wild type or HOIL-1 C460A (Figure S1A). Cells expressing HOIL-1 C460A displayed a pronounced accumulation of high-molecular-weight α-Synuclein species in the insoluble pellet fraction. This was consistent with the formation of large amyloid structures. LUBAC subunits, HOIP, HOIL-1, and SHARPIN, were also enriched in this fraction, suggesting that LUBAC is recruited to or retained within α-Synuclein aggregates (Figure S1A).

Next, we investigated whether the loss of HOIL-1 catalytic activity in an endogenous protein produces similar effects. Mouse embryonic fibroblasts (MEFs) derived from *Hoil-1^C458A/C458A^* mice generated by CRISPR/Ca s9 (Rodriguez Carvajal *et al*., 2021) or *Hoil-1^+/+^*littermate controls were used to assess the physiological relevance of HOIL-1 catalytic activity in an endogenous, non-neuronal context and to confirm that the observed phenotypes were not cell-type specific. We examined the total α-Synuclein and its disease-associated phosphorylated form (pS129) in MEFs (Oueslati, 2016). *Hoil-1^C458A/C458A^*MEFs accumulated larger amounts of α-Synuclein-positive amyloid than *Hoil-1^+/+^*MEFs (Figure S1B, C). Consistent with the pathological aggregation, *Hoil-1^C458A/C458A^*MEFs exhibited prominent pS129-α-Synuclein aggregates, whereas *Hoil-1^+/+^* MEFs displayed minimal staining (Figure S1B, D). These findings demonstrate that the loss of HOIL-1 catalytic activity drives the buildup of both total and pathologically modified α-Synuclein aggregates.

### Aggregate clearance is defective in MEFs expressing catalytically inactive HOIL-1 mutant

To examine how HOIL-1 catalytic activity contributes to proteostasis, we challenged *Hoil-1^+/+^* and *Hoil-1^C458A/C458A^*MEFs with either heat stress (42°C) or lysosomal inhibition using a specific inhibitor of vacuolar H+-ATPase, bafilomycin A1. The aggregates were visualized using Proteostat^®^, a dye that selectively labels misfolded protein aggregates. Under both stress conditions, *Hoil-1^C458A/C458A^* MEFs accumulated significantly larger aggregates than *Hoil-1^+/+^* MEFs (Figure S2A–C), suggesting that the loss of HOIL-1 catalytic activity sensitizes cells to proteotoxic stress.

To determine whether HOIL-1 regulates aggregate formation or clearance, we performed a puromycin chase assay. Puromycin, a tRNA mimic, induces the premature termination of nascent polypeptides, generating misfolded proteins that rapidly coalesce into amorphous aggregates (Lelouard *et al*, 2004; Zaffagnini *et al*, 2018). MEFs were treated with puromycin for 2 h to induce aggregate formation, followed by a 2 h recovery period in puromycin-free medium to allow aggregate clearance (Figure 2A). Aggregates were monitored by immunocytochemistry staining for endogenous p62, a ubiquitin-dependent receptor for misfolded proteins, and ubiquitin. Following puromycin treatment, *Hoil-1^C458A/C458A^* MEFs displayed an increased number of p62-positive puncta (p62 bodies) per cell relative to *Hoil-1^+/+^*MEFs, although their sizes were comparable at this stage (Figure 2B-D). In contrast, differences were observed during the recovery phase. While *Hoil-1^+/+^* MEFs efficiently cleared puromycin-induced p62 bodies, *Hoil-1^C458A/C458A^* MEFs failed to do so and instead accumulated large, irregular, ubiquitin- and p62-positive structures (Figure 2B, E, and F). These findings indicate that although HOIL-1 catalytic activity is dispensable for the initial formation of puromycin-induced aggregates, it is essential for their subsequent clearance.

**Figure 2.**
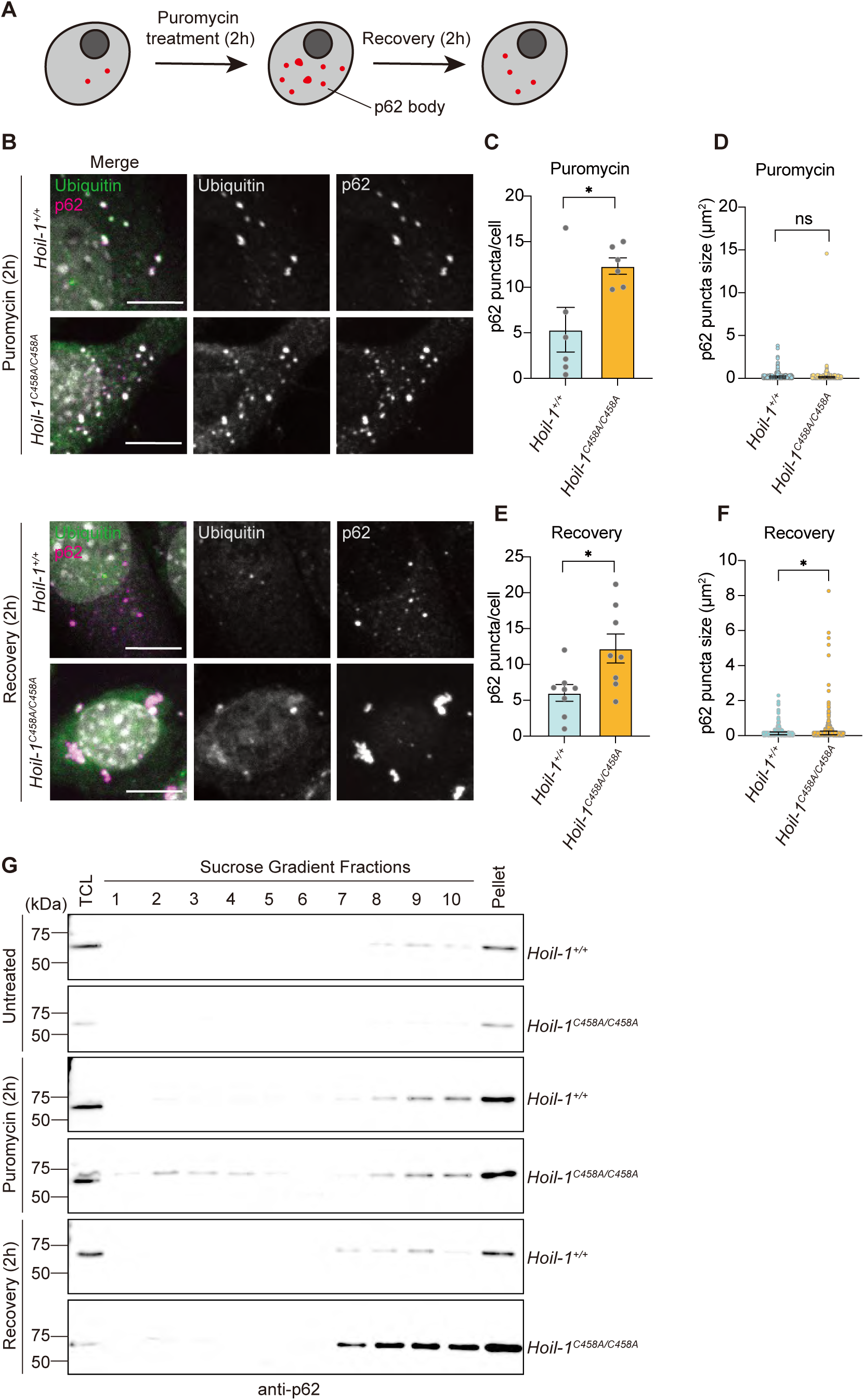
Defect in p62 body clearance in *Hoil-1^C458A/C458A^* cells. A. Schematic of the puromycin chase assay. B. *Hoil-1^+/+^*and *Hoil-1^C458A/C458A^*MEFs were treated with 5 μg/ml puromycin for 2 h and placed in puromycin-free media for 2 h for recovery. Representative images showing the colocalization of ubiquitin and p62 after puromycin treatment and 2 h after recovery in *Hoil-1^+/+^*and *Hoil-1^C458A/C458A^* MEFs. Scale bar = 5 μm. Representative data from three independent biological replicates. C, D. Graphical representation of (C) puncta/cell and (D) p62 puncta sizes after treatment with 5 μg/ml puromycin for 2 h. (C) Mean ± SEM was plotted, and unpaired t-test statistical analyses were performed; *, P=0.0235. (D) The median with interquartile range was plotted, and the non-parametric Mann-Whitney test was performed; ns, P>0.05. N=3 independent biological repeats (puromycin treatment, *n*= 68 *Hoil-1^+/+^*cells and *n*= 38 *Hoil-1^C458A/C458A^* cells; recovery, *n*= 48 *Hoil-1^+/+^* cells and *n*= 60 *Hoil-1^C458A/C458A^*cells). E, F. Graphical representation of (E) puncta/cell and (F) p62 puncta sizes under recovery conditions for 2 h. (E) Mean ± SEM was plotted, and unpaired t-test statistical analyses were performed; *, P=0.019. (F) Median with interquartile range was plotted and non-parametric Mann-Whitney test was performed; *, P=0.0315. N=3 independent biological repeats (puromycin treatment, *n*= 68 *Hoil-1^+/+^* cells and *n*= 38 *Hoil-1^C458A/C458A^* cells; recovery, *n*= 48 *Hoil-1^+/+^* cells and *n*= 60 *Hoil-1^C458A/C458A^*cells). G. Sucrose gradient fractionation to compare the distribution of p62 in *Hoil-1^+/+^* and *Hoil-1^C458A/C458A^* MEFs under untreated, 5 mg/ml puromycin (2 h) treatment, and recovery (2 h) conditions as indicated. Representative data are from N=3 independent repeats.

To verify that p62 bodies corresponded to insoluble aggregates, we performed sucrose gradient fractionation of lysates from MEFs subjected to puromycin treatment and recovery. Consistent with microscopic observations, denser fractions from *Hoil-1^C458A/C458A^* MEFs contained increased signals of p62 specifically during the recovery phase (Figure 2G). Together, these results demonstrate that the loss of HOIL-1 catalytic activity impairs the efficient clearance of misfolded protein aggregates positive for p62 and ubiquitin, leading to their aberrant accumulation.

### Aggrephagy-mediated clearance of p62 bodies is compromised in *Hoil-1^C458A/C458A^*MEFs

Protein aggregates are removed primarily through selective autophagy called aggrephagy (Bauer *et al*., 2023; Ichimura *et al*, 2008). Because *Hoil-1^C458A/C458A^*MEFs accumulated p62^+^/ubiquitin^+^ puncta (Figure 2B) and Proteostat®-stained aggregates (Figure S2A-C) under proteotoxic stress, we investigated whether impaired aggrephagy underlies this defect. We first examined the recruitment of key autophagy machinery to p62 bodies under proteotoxic stress using puromycin treatment and subsequent recovery. In both *Hoil-1^+/+^* and *Hoil-1^C458A/C458A^*MEFs, p62 puncta colocalized with LC3, indicating that autophagy initiation and autophagosome assembly at p62 bodies remained intact. Consistent with earlier observations, *Hoil-1^C458A/C458A^* MEFs displayed larger p62 puncta, but the overall percentage of p62–LC3 overlap was comparable between the genotypes in both the puromycin and recovery phases (Figure 3A–C).

**Figure 3.**
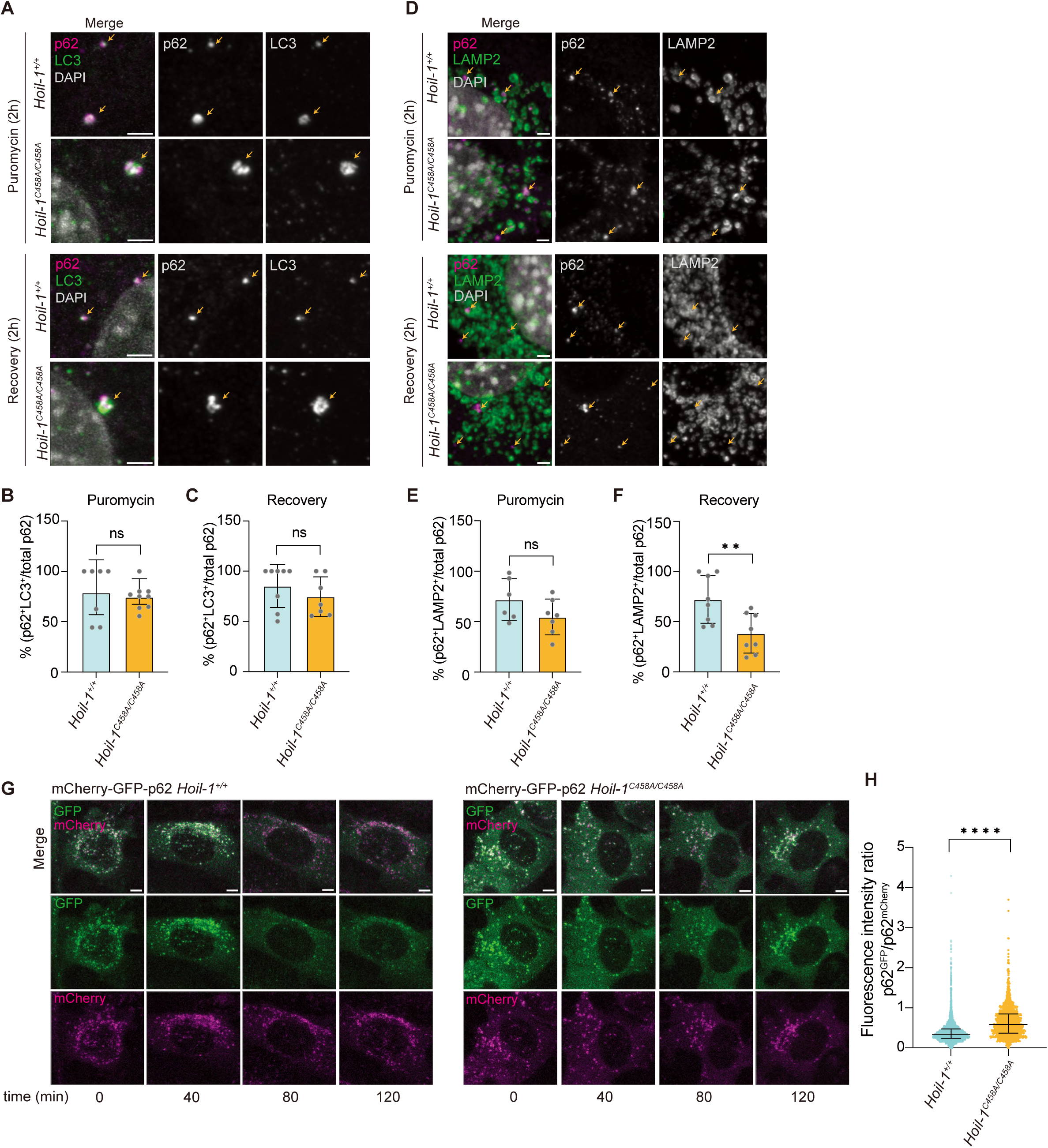
Aggrephagy-mediated clearance of aggregates is compromised in *Hoil-_1C458A/C458A_* _MEFs_. A. Colocalization of p62 and LC3 in *Hoil-1^+/+^* and *Hoil-1^C458A/C458A^* MEFs upon 5 μg/ml puromycin treatment and recovery as indicated. Scale bar = 5 μm. B-C. Graphical representation of the percentage of p62 colocalized with LC3 after 5 μg/ml puromycin treatment (B) and after recovery (C). Values plotted are mean ± SEM from N=2 independent biological repeats (puromycin, *Hoil-1^+/+^* n= 7 fields and *Hoil-1^C458A/C458A^* n= 8 fields: recovery, *Hoil-1^+/+^*n= 8 fields and *Hoil-1^C458A/C458A^* n= 8 fields). Unpaired t-test statistical analysis was performed (ns, P > 0.05). D. Colocalization of p62 and LAMP2 in *Hoil-1^+/+^* and *Hoil-1^C458A/C458A^* MEFs upon 5 μg/ml puromycin treatment (2 h) and recovery (2 h) as indicated. Scale bar = 5 μm. E-F. Graphical representation of the percentage of p62 colocalizing with LAMP2 protein after5 μg/ml puromycin treatment (E) and recovery (F). Values plotted are mean ± SEM from N=3 independent biological repeats (puromycin, *Hoil-1^+/+^* cells n= 6 fields and *Hoil-1^C458A/C458A^* cells n= 7 fields; recovery, n= 7 fields from *Hoil-1^+/+^* and *Hoil-1^C458A/C458A^* MEF samples). Unpaired t-test statistical analysis was performed; ns, P>0.05; **, P=0.0078. G. Representative images of time-lapse imaging to monitor autophagy flux of GFP-mCherry-p62 *Hoil-1^+/+^* and *Hoil-1^C458A/C458A^* MEFs under puromycin treatment (*n*= 40 cells from N=3 independent biological repeats). Scale bar = 5 μm. H. Graphical representation of the ratio of p62^GFP^/ p62^mCherry^ fluorescence intensities of mCherry-GFP-p62 stably expressing *Hoil-1^+/+^* and *Hoil-1^C458A/C458A^* MEFs treated with 5 μg/ml puromycin. Values plotted are median fluorescence intensity ratios with interquartile range of *n*=40 cells from N=3 independent biological repeats. Mann-Whitney statistical analysis was performed (**** P<0.0001).

To evaluate whether p62 bodies are degraded by lysosomes, we examined the colocalization of p62 bodies with LAMP2, a lysosomal membrane marker. Following puromycin treatment, p62-bodies in both *Hoil-1^+/+^* and *Hoil-1^C458A/C458A^* MEFs were surrounded by LAMP2-positive structures (Figure 3D, E, and Figure S3A, B). During recovery, *Hoil-1^+/+^* MEFs showed efficient recruitment of LAMP2 to p62 puncta and marked reduction in p62-body size. In contrast, *Hoil-1^C458A/C458A^* MEFs accumulated enlarged p62 bodies that lacked adjacent LAMP2-positive structures, indicating failure of autophagosome–lysosome fusion or autolysosome maturation (Figures 3D, F, and Figure S3B).

To directly monitor the p62-dependent autophagic flux, we generated reporter MEF lines stably expressing GFP–mCherry–p62 (Pankiv *et al*, 2007). In this system, GFP fluorescence is quenched under acidic conditions, such as in lysosomes, whereas mCherry fluorescence remains stable, allowing distinction between autophagosomes (GFP^+^ mCherry^+^) and autolysosomes (GFP^−^mCherry^+^). In the *Hoil-1^+/+^* reporter line, puromycin treatment triggered a progressive loss of GFP signal, indicating that p62 bodies were delivered to lysosomes. In contrast, *Hoil-1^C458A/C458A^* MEFs accumulated GFP^+^ mCherry^+^ p62 puncta, indicating that lysosomal degradation was blocked (Figure 3G). Quantification of the GFP/mCherry ratios confirmed significantly elevated levels of GFP retention in *Hoil-1^C458A/C458A^*reporter MEFs (Figure 3H).

To further validate these findings, we generated GFP–mCherry–LC3 reporter MEFs (Ebner *et al*, 2018; Kimura *et al*, 2007). Consistent with the p62 reporter results, *Hoil-1^C458A/C458A^* MEFs treated with puromycin accumulated dual-positive (GFP^+^ mCherry^+^) LC3 vesicles, indicating defective autophagic flux and impaired autophagosome maturation specifically during aggregate clearance (Figure S3C and D).

Together, these data suggest that although early autophagy components, such as LC3, are effectively recruited to p62 bodies, the late stages of aggrephagy, particularly autolysosome formation at the site, are compromised in *Hoil-1^C458A/C458A^* MEFs. Importantly, the mRNA level of autophagy-related genes was comparable between *Hoil-1^+/+^* and *Hoil-1^C458A/C458A^* MEFs (Figure S3E). Thus, HOIL-1 catalytic activity appears to be dispensable for general autophagy but is specifically required for efficient aggregate-selective autophagy.

### Transition in condensate properties in *Hoil-1^C458A/C458A^* MEFs

To determine whether the altered biophysical states of p62-positive aggregates contribute to clearance defects in *Hoil-1^C458A/C458A^*MEFs, we first assessed the circularity of endogenous p62 puncta (Ray *et al*, 2020; Tomaszewski *et al*, 2023). Soluble condensates (also termed liquid-like biomolecular condensates) typically display near-perfect circularity (close to 1), whereas solid aggregates exhibit irregular and non-circular shapes. Stratification of p62 puncta by size in the recovery samples of *Hoil-1^C458A/C458A^* MEFs revealed a progressive decrease in circularity with increasing puncta size, consistent with a shift toward more rigid and solid-like structures (Figure S4A).

To evaluate whether these structures represent solid aggregates, we examined their sensitivity to 1,6-hexanediol (1,6-HD), a chemical disruptor of weak hydrophobic interactions (Yamazaki *et al*, 2018). As a positive control, FUS R495X, which forms 1,6-HD -sensitive condensates in human embryonic kidney 293T (HEK293T) cells, was readily dispersed by 1,6-HD (Figure S4B). In *Hoil-1^+/+^*MEFs, puromycin-induced p62^+^ ubiquitin^+^ puncta were efficiently dissolved by 1,6-HD. However, in *Hoil-1^C458A/C458A^* MEFs, a substantial fraction of p62 puncta persisted after treatment, indicating reduced 1,6-HD sensitivity and supporting a transition to more solid-like structures (Figure 4A, B).

**Figure 4.**
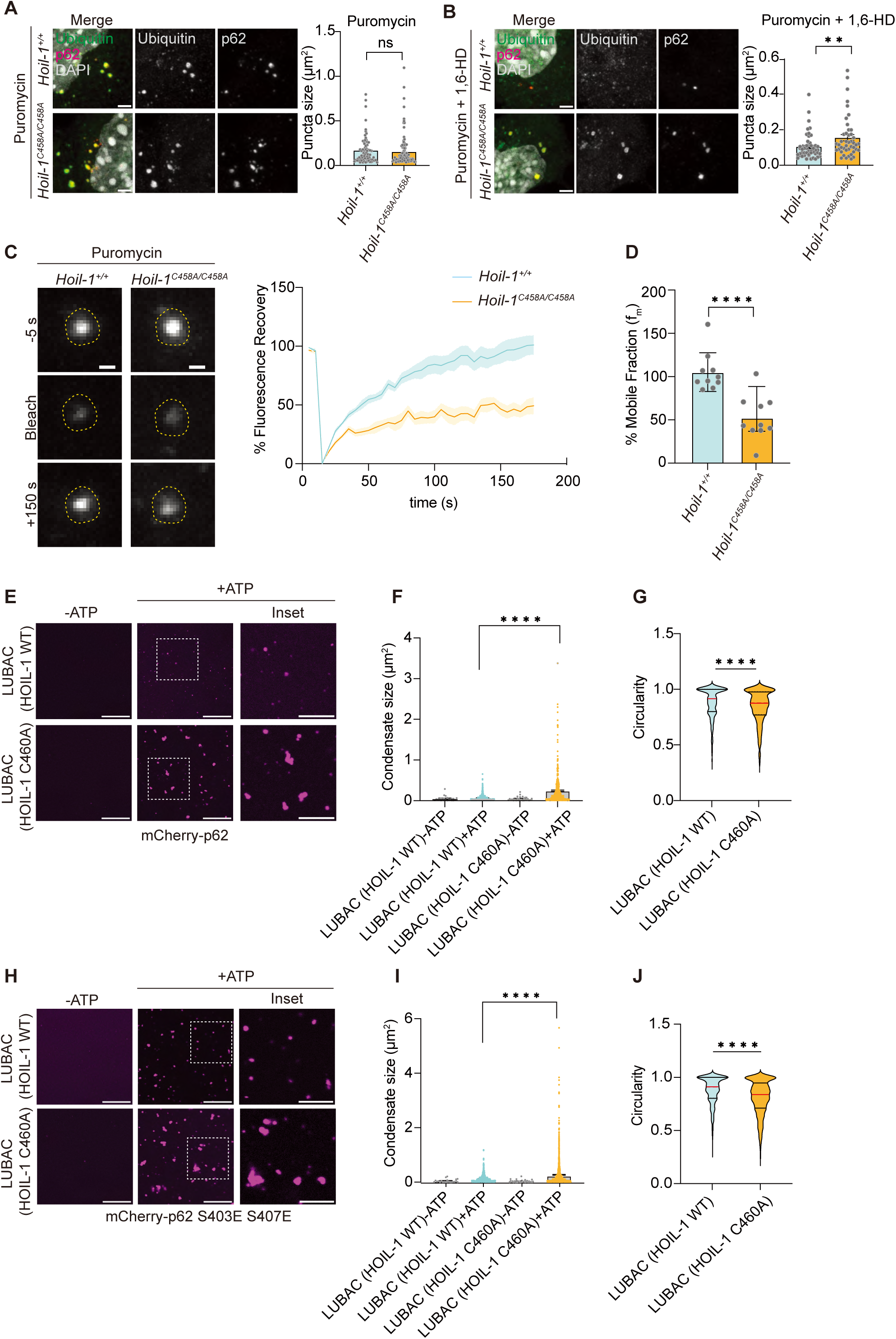
Transition of condensate property by catalytically inactive mutations in HOIL-1 in MEFs and *in vitro*. A-B. Representative images of immunofluorescence of ubiquitin (green) and p62 (red) proteins, under (A) 5 μg/ml puromycin treatment for 2 h and (B) 5 μg/ml puromycin (2 h) + 5% 1,6-Hexanediol (1,6-HD), to demonstrate the dissolution of puromycin-induced p62 protein condensates under 5% 1,6-HD treatment in *Hoil-1^+/+^* and *Hoil-1^C458A/C458A^* MEFs. Representative data from N=3 biological replicates. Scale bar = 2 μm. Graphical representation of p62 aggregate size under (A) puromycin treatment and (B) puromycin + 5% 1,6-HD treatment in *Hoil-1^+/+^*and *Hoil-1^C458A/C458A^* MEFs. Values plotted are mean ± SEM from N=3 independent biological repeats (puromycin, *Hoil-1^+/+^* cells n= 70 and *Hoil-1^C458A/C458A^*cells *n*= 97; recovery, *Hoil-1^+/+^* cells *n*= 51 and *Hoil-1^C458A/C458A^* cells *n*= 44). Mann-Whitney statistical analysis was performed; ns, P>0.05, **, 0.0071. C. Representative images of Fluorescence Recovery After Photobleaching (FRAP) analysis in GFP-p62 expressing *Hoil-1^+/+^* and *Hoil-1^C458A/C458A^*MEFs at pre-bleaching, bleached, and 150 s post-recovery. A graphical representation of the mean % fluorescence recovery ± SEM with respect to time (s) is shown. Data from *n*= 9 events from N= 3 independent biological repeats. Scale bar = 0.5 μm. D. Graphical representation of the mean mobile fractions (f_m_) ± SD of GFP-p62 between *Hoil-1^+/+^* and *Hoil-1^C458A/C458A^* MEFs determined by FRAP analysis. Statistical analysis was performed using an unpaired t-test, **** P<0.0001. N= 3. E. Representative fluorescence images of *in vitro* LLPS assay with mCherry-p62 WT and Met1-linked ubiquitin chains generated by LUBAC WT (LUBAC (HOIL-1 WT)) or LUBAC containing HOIL-1 C460A (LUBAC (HOIL-1 C460A)). F, G. Graphical representation of (F) condensate size and (G) circularity of mCherry-p62 condensates obtained from the LLPS assay (E). (F) Data plotted are the mean ± SEM; statistical analysis was performed using the Kruskal-Wallis test with Dunn’s multiple comparisons, **** P<0.0001. (G) Data plotted are median (red line) and quartiles (black lines); statistical analysis was performed using non-parametric Mann-Whitney test, **** P<0.0001. N=3; scale bar = 10 μm, inset = 5 μm. H. Representative fluorescence images of the *in vitro* LLPS assay with mCherry-p62 S403E S407E and Met1-linked ubiquitin chains generated by LUBAC WT (LUBAC (HOIL-1 WT)) or LUBAC containing HOIL-1 C460A (LUBAC (HOIL-1 C460A)). I, J. Graphical representation of (I) condensate size and (J) circularity of the mCherry-p62 S403E S407E condensates obtained from the LLPS assay (H). (I) Data plotted are the mean ± SEM; statistical analysis was performed using the Kruskal-Wallis test with Dunn’s multiple comparison ****, P<0.0001. (J) Data plotted are median (red line) and quartiles (black lines); statistical analysis was performed using the non-parametric Mann-Whitney test, **** P<0.0001. N=3; scale bar = 10 μm, inset = 5 μm.

Next, we evaluated the dynamic properties of these condensates using fluorescence recovery after photobleaching (FRAP). In puromycin-treated MEFs, in which GFP–p62 was stably expressed, GFP^+^ puncta showed robust fluorescence recovery in *Hoil-1^+/+^* MEFs, consistent with rapid internal molecular exchange. In contrast, GFP–p62 puncta in *Hoil-1^C458A/C458A^* MEFs showed markedly slower and incomplete recovery, indicative of reduced molecular mobility and a shift toward a less dynamic, solid state (Figure 4C, D).

Our previous work demonstrated that, *in vitro*, LUBAC containing the HOIL-1 C460A catalytically inactive mutant produces Met1-linked ubiquitin chains more efficiently than wild-type LUBAC (Rodriguez Carvajal *et al*., 2021). This altered activity leads to the depletion of short ubiquitin chains and the absence of oxyester-linked ubiquitin species, which are typically generated by wild-type LUBAC. Given that LUBAC containing the HOIL-1 C460A mutant produces aberrant Met1-linked ubiquitin chains *in vitro*, we investigated how these chains influence the aggregate properties. We utilized Met1-linked chains produced *in vitro* by either HOIL-1 wild type or HOIL-1 C460A-containing LUBAC (Figure S4C) in liquid-liquid phase separation (LLPS) reconstitution assays using recombinant mCherry-p62 (Figure 4E-J). Met1-linked chains synthesized by HOIL-1 C460A-containing LUBAC promoted the formation of large, irregular, non-spherical p62 condensates, in contrast to the smaller, more rounded condensates formed by LUBAC wild type-derived ubiquitin chains (Figure 4E-G). Interestingly, p62 with phosphomimetic mutations (S403E/S407E) in its UBA domain, known for its enhanced ubiquitin affinity(Lim *et al*, 2015)⍰, formed large, irregular condensates in the presence of ubiquitin chains generated by LUBAC containing HOIL-1 C460A (Figure 4H-J). These aberrant condensates were similar to those observed in a reaction using wild-type p62. These aberrant condensates closely resembled the solid aggregates observed in *Hoil-1^C458A/C458A^* ⍰

Collectively, these findings indicate that HOIL-1 catalytic activity plays a crucial role in maintaining the proper physical state of the Met1-linked ubiquitin-dependent p62 condensates.

### Elevated Met1-linked ubiquitin chains contribute to solid-like aggregate formation

Our previous findings indicated that the levels of Met1-linked ubiquitin chains were elevated in *Hoil-1^C458A/C458A^* MEFs upon TNF stimulation (Rodriguez Carvajal *et al*., 2021). To determine whether increased levels of Met1-linked ubiquitin chains contribute to aggregate formation, we enriched Met1-linked chains from *Hoil-1^+/+^*and *Hoil-1^C458A/C458A^* MEFs treated with puromycin using GST-tagged triplicates of the UBAN domain of ABIN-1, which specifically binds to Met1-linked ubiquitin chains (Fennell *et al*, 2020). *Hoil-1^C458A/C458A^* MEFs accumulated more Met1-linked chains upon puromycin treatment than *Hoil-1^+/+^*MEFs, indicating that Met1-linked ubiquitin chains were more efficiently generated in HOIL-1 catalytic mutant cells under proteotoxic stress (Figure 5A).

**Figure 5.**
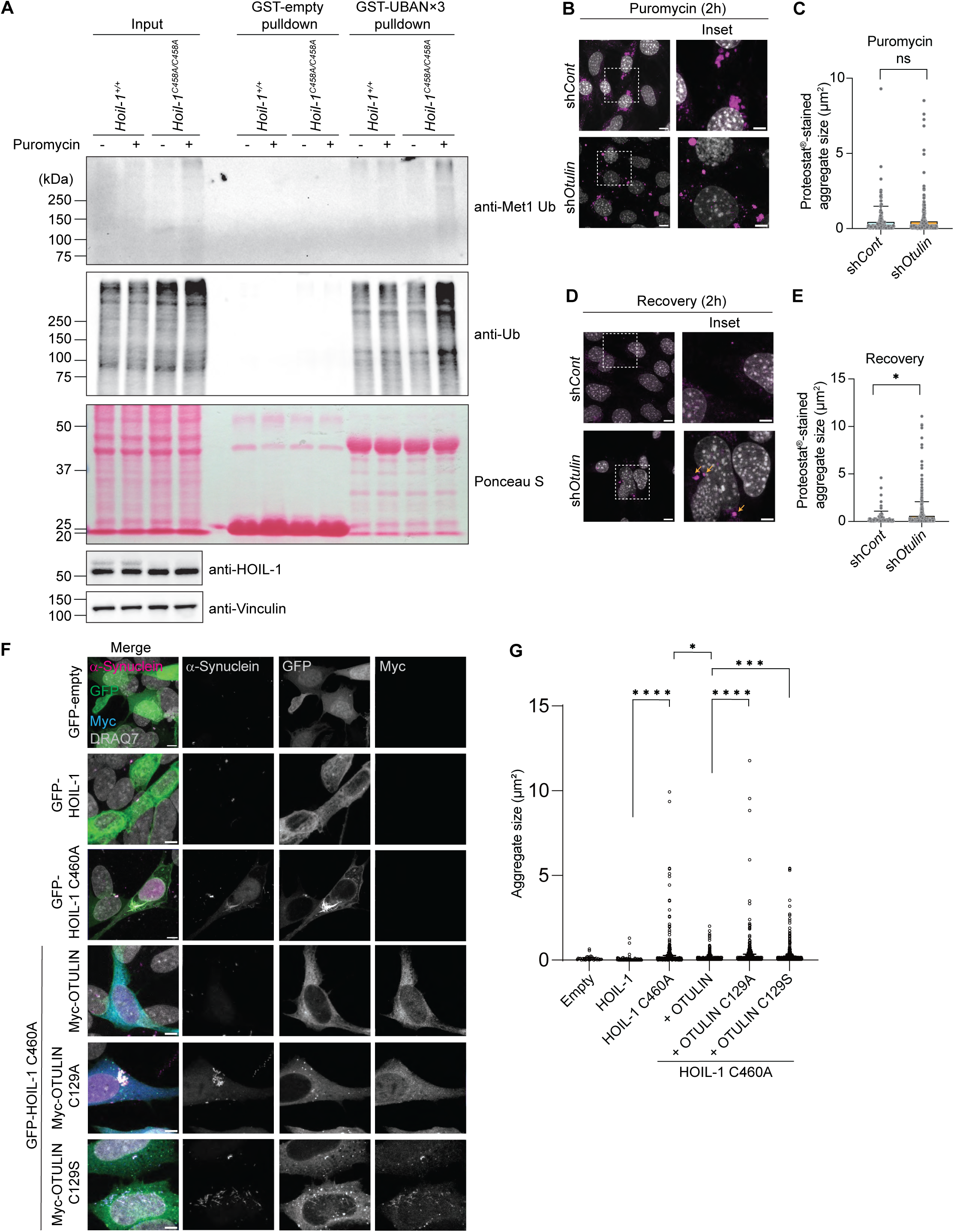
Accumulation of Met1-linked ubiquitin chains upon proteotoxic stress in Hoil-1^C458A/C458A^ MEFs. A. GST-empty and GST-UBAN×3 pulldown from *Hoil-1^+/+^* and *Hoil-1^C458A/C458A^* MEF lysates followed by immunoblotting analysis. Ponceau S (bottom panel) was used as the loading control. Representative data from N=3 independent biological repeats. B. Comparison of Proteostat^®^-stained aggregates in sh*Cont* and sh*Otulin* MEFs treated with 5 μg/ml puromycin. Representative images from N=3 independent biological repeats. Scale bar = 5 μm. C. Graphical representation of aggregate size under puromycin treatment in sh*Cont* and sh*Otulin* MEFs. Data plotted are mean ± SEM; Mann-Whitney statistical analysis was performed; ns, P>0.05; >75 cells were counted for each set from N=3 independent biological repeats. D. Comparison of Proteostat^®^-stained aggregates in sh*Cont* and sh*Otulin* MEFs under recovery conditions. Representative images from N=3 biological replicates. Scale bar = 5 μm. E. Graphical representation of aggregate size under the recovery conditions in sh*Cont* and sh*Otulin* MEFs. Data plotted are mean ± SEM; Mann-Whitney statistical analysis was performed; *, P= 0.023; >75 cells were counted for each set from N=3 independent biological replicates. F. Representative microscopy images to show the colocalization of α-Synuclein aggregates and GFP-tagged HOIL-1 or HOIL-1 C460A, co-expressed with Myc-tagged OTULIN, OTULIN C129A or OTULIN C129S as indicated, in SH-SY5Y cells. Representative images from N=2 biological repeats. Scale bar = 5 μm. G. Graphical representation of the aggregate mean sizes ± SEM from (F). Statistical analysis was performed using the Kruskal-Wallis test with Dunn’s multiple comparisons; *, P= 0.0311, ***, P= 0.0002, ****, P<0.0001; >250 cells were counted for each set from N=2 independent biological repeats.

Next, we stably knocked down OTULIN, a deubiquitinase specific for Met1-linked ubiquitin chains, in MEFs (Figure S5A) and performed GST pulldown following puromycin treatment. Although basal Met1-linked chains were already elevated in sh*Otulin* MEFs, we observed a modest increase in the signal of Met1-linked ubiquitin chains after puromycin treatment; nevertheless, the enriched Met1-linked chains were clearly higher than those in control MEFs (Figure S5B). Supporting our hypothesis that elevated Met1-linked ubiquitin chains sensitize cells to form larger aggregates, sh*Otulin* MEFs developed significantly larger p62-positive puncta during the puromycin recovery phase and under heat stress, mirroring the phenotype observed in *Hoil-1^C458A/C458A^*MEFs (Figure 5B–E, S5C–E).

To further test whether excessive Met1-linked chains are a major driver of aggregate accumulation, we examined whether OTULIN overexpression could ameliorate the effects induced by the HOIL-1 catalytically inactive mutant (Figure 5F, G). To this end, we co-expressed GFP, GFP-HOIL-1, or GFP-HOIL-1 C460A with α-Synuclein in SH-SY5Y cells (Figure 5F). Consistent with our earlier observations, α-Synuclein aggregates in HOIL-1 C460A–expressing cells were significantly larger than those in cells expressing wild-type HOIL-1 (Figure 5G). In the HOIL-1 C460A background, we overexpressed wild-type OTULIN or its catalytically inactive mutants (OTULIN C129A or C129S). This allowed us to assess whether reducing the excess Met1-linked chains could diminish the size of the aggregates formed in these cells. We observed a significant reduction in α-Synuclein aggregate size upon OTULIN overexpression, emphasizing the contribution of Met1-linked chains to driving aggregate accumulation.

In summary, these results support a model in which the excessive expression of Met1-linked ubiquitin chains in cells promotes the formation of more solid and less dynamic protein assemblies, potentially contributing to impaired aggregate clearance (**Figure 6**).

**Figure 6.**
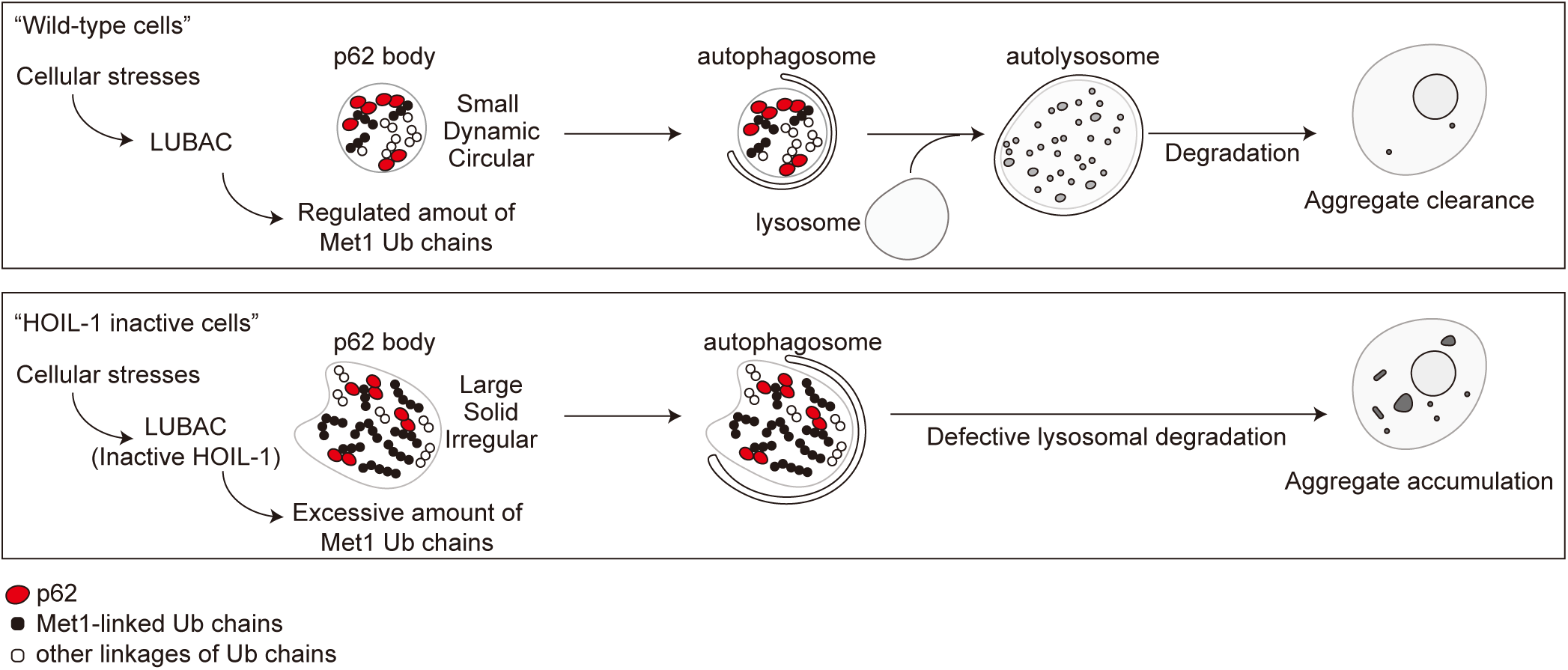
Mechanism of abrogated protein aggregate clearance in the presence of *Hoil-1^C458A/C458A^* catalytically inactive mutant. Under proteotoxic stress, normal cells tag misfolded proteins with ubiquitin chains, including the oxyester-branched Met1-linked ubiquitin chains generated by LUBAC. These ubiquitinated substrates are then concentrated in p62 bodies, which function as dynamic, liquid-like condensates. p62 bodies are subsequently processed through the aggrephagy pathway, which involves the recruitment of autophagosomes, maturation through lysosomal fusion to form autolysosomes, and eventual degradation and recycling of sequestered material. This sequence of events maintains cellular proteostasis by ensuring efficient clearance of stress-induced aggregates. However, in cells expressing the HOIL-1 catalytically inactive mutant, excess Met1-linked ubiquitin chains accumulate because of unrestrained HOIP activity. p62 bodies in this context become overloaded with Met1-linked chains, shifting from a dynamic, liquid-like state to a more rigid, solid-like architecture. These less dynamic aggregates are inefficiently processed by aggrephagy and persist as insoluble deposits within the cell.

## Discussion

Our findings underscore the importance of a precise balance between ubiquitination and deubiquitination of Met1-linked ubiquitin chains in cellular proteostasis. Specifically, the catalytic activity of HOIL-1 is critical for fine-tuning the function of HOIP within LUBAC, thereby preventing overproduction of Met1-linked ubiquitin chains. This regulatory mechanism ensures that in the presence of functional HOIL-1, the overall quantity and potentially the architecture of Met1-linked ubiquitin chains are tightly controlled. We found that elevation of Met1-linked chains, either through HOIL-1 catalytic inactivation or knockdown of the Met1-linked chain-specific deubiquitinase OTULIN, led to defects in aggregate clearance. This suggests that excess Met1-linked ubiquitin may actively contribute to aggregate stabilization and hinder their resolution. Consistent with these cellular observations, our *in vitro* LLPS assay suggests that excessive Met1-linked chains may promote the formation of rigid, solid-like aggregates, a phenomenon directly supported by our cellular biophysical analyses (e.g., 1,6-HD sensitivity and FRAP). These observations were consistent across neuronal cells and MEFs, suggesting a conserved role oof HOIL-1 in aggregate regulation. Consistent with previous studies (Furthmann *et al*., 2023; van Well *et al*., 2019; Zhang *et al*, 2022), we found that LUBAC components were sequestered in aggregates, as evidenced by microscopy and gradient fractionation of soluble and insoluble proteins, confirming the direct involvement of LUBAC in aggregate processing.

Our study implicates selective autophagy, namely aggrephagy, as the primary pathway affected by HOIL-1 loss-of-function. While early autophagy steps remained intact, as evidenced by p62-LC3 colocalization, late-stage autophagy was critically compromised in *Hoil-1^C458A/C458A^* MEFs, highlighting a defect in autolysosome formation and, thus, aggrephagy-mediated degradation of protein aggregates. This impaired clearance is likely exacerbated by the observed transition of p62 bodies from dynamic, liquid-like condensates to more rigid, solid-like structures in the absence of functional HOIL-1. Given the importance of liquidity in early aggregate sequestration and efficient autophagic clearance (Komatsu, 2022; Kurusu *et al*, 2024; Turco *et al*, 2021; Yamasaki *et al*, 2020), our observations suggest that HOIL-1 catalytic activity is essential for maintaining the proper biophysical properties of aggregates, thereby facilitating their efficient turnover.

The precise molecular mechanism by which excessive Met1-linked ubiquitin chains shape the biophysical properties of aggregate materials remains unclear. One possibility is that HOIP directly extends Met1-linked chains on pre-existing ubiquitinated substrates within aggregates, altering their surface properties or internal structures. Another possibility is that free Met1-linked chains are incorporated into growing p62 bodies, thereby directly influencing their physical behavior and promoting solidification. Indeed, p62 and all three components of LUBAC (HOIL-1, HOIP and SHARPIN) have ubiquitin interaction domains raising the possibility of interaction between ubiquitin chains and these molecules within p62 bodies. However, these mechanisms are not mutually exclusive and require further investigation. Our findings suggest that excess Met1-linked ubiquitin chains lower the threshold for phase transitions, promoting more rigid aggregate states when HOIL-1-mediated catalytic restraint is lost. Although our study provides compelling evidence for the role of HOIL-1 catalytic activity in proteostasis, several important future directions should be considered. The current study primarily relied on *in vitro* and *in cellulo* models; therefore, future studies should aim to validate these findings in relevant *in vivo* neurodegeneration models to confirm the physiological impact of HOIL-1 deficiency on aggregate clearance and disease progression. Furthermore, identifying the specific substrates within aggregates targeted by LUBAC and characterizing the precise topology (e.g., chain length and branching points) of Met1-linked ubiquitin chains in HOIL-1 mutant cells will be crucial for fully elucidating the mechanism by which they promote aggregate solidification. Finally, investigation of the potential for therapeutic intervention by modulating HOIL-1 activity or Met1-linked chain levels represents a promising avenue for developing novel treatments for neurodegenerative proteinopathies.

## Materials and Methods

### Plasmids

pcDNA 3.1 human HOIL-1, pcDNA 3.1 human HOIL-1 C460A, pEGFP-C1 human HOIL-1, and pEGFP-C1 human HOIL-1 C460A were previously generated (Gomez-Diaz *et al*, 2021). p62 was inserted into the EcoRI/SalI restriction sites of the pBABE-puro-mCherry-EGFP vector using standard subcloning methods. pGex-4T1-mouse ABIN1-UBAŃ3 was generated and used previously (Fennell *et al*., 2020). pRK-Myc-human OTULIN, pRK-Myc-human OTULIN C129A, and pRK-Myc-human OTULIN C129S were generated using standard subcloning methods. pcDNA5/FRT/TO/Flag X3-FUS R495X was a gift from the Hirose Lab (The University of Osaka) (Yamazaki *et al*, 2012). pMXs-puro GFP-mouse p62 (Addgene #38277) and pMXs-IP GFP-human NBR1 (Addgene #38283) were gifts from Noboru Mizushima (Itakura & Mizushima, 2011). pBABE-puro-mCherry-EGFP-human LC3B (Addgene #22418) was a gift from Jayanta Debnath (N’Diaye *et al*, 2009). The pGEX6P-mCherry-human p62 and pGEX6P-mCherry-human p62 S403E S407E plasmids were described previously (Ikeda *et al*, 2023). pLKO.1-shRNA-m*Otulin* (shRNA hairpin seq: GCTCATACTGTTACCAGAGAACTCGAGTTCTCTGGTAACAGTATGAGCT) and pLKO.1- shRNA-*control* scramble shRNA (shRNA hairpin seq: CCTAAGGTTAAGTCGCCCTCGCTCGAGCGAGGGCGACTTAACCTTAGG) were generated by standard subcloning methods.

### Antibodies

Anti-HOIL-1, clone 2E2 (Millipore, MAB8576), anti-human RNF31/HOIP (Aviva Systems Biology, ARP43241_P050), anti-SHARPIN (Novus, NBP2-04116), anti-Vinculin (Sigma, V9131), anti-LC3 (Nanotools, 0260-100/LC3-2G6), anti-LAMP2 (Abcam, ab13524), anti-p62 (MBL, PM045), anti-α-Synuclein [MJFR1] (Abcam, ab138501), anti-phosphorylated α-Synuclein (Wako, 015-25191), and anti-ubiquitin [FK2] (Enzo, BML-PW8810-0500), anti-OTULIN (Abcam, ab151117), anti-c-Myc [9E10] (Covance, MMS-150P), anti-ubiquitin (P4D1) (Santa Cruz Biotechnology, sc-8017), and anti-Met1 ubiquitin [LUB9] (Merck, MABS451) antibodies were used in this study.

### Cell lines

SH-SY5Y cells (ECACC, 94030304) were maintained at 37 °C in 5% CO_2_ and grown in a 1:1 ratio of Ham’s F12 medium (Wako, 087-08335) and EBSS-MEM (Wako, 055-08975) supplemented with 15% fetal bovine serum (FBS) and 100 U/ml penicillin-streptomycin (Nacalai Tesque, 09367-34). Immortalized MEFs derived from *Hoil-1^+/+^* and *Hoil-1^C458A/C458A^*mice were established as described previously (Rodriguez Carvajal *et al*., 2021). Human embryonic kidney 293T (HEK293T) cells (ATCC) and the packaging cell line Platinum-E (Plat-E) (Markowitz *et al*, 1990; Morita *et al*, 2000) were maintained at 37 °C in 5% CO_2_ in Dulbecco’s modified Eagle’s medium high glucose (Wako, 044-29765) supplemented with 10% fetal calf serum (Biowest, S1810-500) and 100 U/ml penicillin-streptomycin (Nacalai Tesque, 09367-34).

### Retroviral and lentiviral infection

Retrovirus production was performed as described previously (Ikeda *et al*, 2007). Briefly, the retroviral plasmids pBABE-puro-mCherry-EGFP-p62, pMXs-puro GFP-p62, and pBABE-puro-mCherry-EGFP-LC3B were transfected into Plat-E cells. At 48 h post-transfection, MEFs were transduced with filtered conditioned media containing retroviral particles using a 0.45 μm syringe PVDF filter unit (Millipore, SLHVR33RS) supplemented with 8 µg/ml polybrene (Sigma-Aldrich, H9268). To obtain stable cell lines, cells were selected using 4 µg/ml puromycin (ROTH, 0240.3). Lentiviral production was performed as described previously (Kumari *et al*, 2014). Briefly, lentiviral plasmids (pLKO.1-shOtulin or pLKO.1-shControl) were transfected into HEK293T cells along with the psPAX2 packaging plasmid and the pMD2.G envelope plasmid. At 36 h post-transfection, MEFs were transduced with filtered conditioned media containing lentiviral particles supplemented with 4 µg/ml polybrene (Sigma-Aldrich, H9268). To obtain stable cell lines, cells were selected using 4 µg/ml puromycin (ROTH, 0240.3).

### Western blotting analysis

This method has been previously described (Ikeda *et al*., 2011). Cells were lysed in chilled lysis buffer (50 mM HEPES, 150 mM NaCl, 1 mM EDTA, 1 mM EGTA, 1% Triton X-100, 10% glycerol, 25 mM NaF, 10 mM ZnCl2, 10 mM NEM, 1 mM PMSF, 5 mM Na3VO4, pH 7.4 supplemented with cOmplete Mini EDTA-free protease inhibitor cocktail (Roche, 11836170001)), centrifuged at 10,000 RCF to remove cell debris, and total cell lysates were resolved by SDS-PAGE, and transferred onto a nitrocellulose (Cytiva, 10600002 (0.45 µm pore size), 10600011 (0.2 µm pore size)) or polyvinylidene difluoride membrane (Millipore, ISEQ00010). Membranes were blocked in 5% <u>bovine serum albumin</u> (BSA)-TBS and blotted with indicated antibodies in blocking solution at 4 °C overnight. HRP-conjugated secondary antibodies were used: goat anti-mouse HRP (BioRad, 170–6516, Cell Signalling Technologies, 7076S) and goat anti-rabbit HRP (Dako, P0448). Western blotting Luminol Reagent (Santa Cruz, sc-2048) and FUSION SOLO S chemiluminescence detection system (Vilber) were used for capturing the chemiluminescent signals.

### Aggregate formation in cells

Aggregates were formed in SH-SY5Y cells grown on coverslips following the manufacturer’s instructions to either create ⍺-Synuclein aggregates (⍺-Synuclein Aggregation Assay Kit, Cosmo Bio, CSR-SYN01) or tau aggregates (Tau Aggregation Assay Kit, Cosmo Bio, TAU01). Cells were simultaneously transfected with pEGFP-C1 human HOIL-1 or pEGFP-C1 human HOIL-1 C460A plasmids. The cells were grown for 72 h after transfection. Aβ (1-42) (Bachem, 4014447) peptide was solubilized as described previously (Stine *et al*, 2003). Oligomeric Aβ was prepared by dissolving Aβ in DMSO and phenol-free F-12 to a final concentration of 100 μM. The mixture was vortexed and stored at 4 °C for 24 h. The next day, SH-SY5Y cells were transfected with pEGFP-C1 human HOIL-1 or pEGFP-C1 human HOIL-1 C460A and treated with 20 μM Aβ for 24 h. Aggregates were detected using the Amyloid Fluorescent Staining Kit (Cosmo Bio, CSR-SYN02) according to the manufacturer’s instructions, except that the nuclei were counterstained with DRAQ7 (Abcam, D15106). Coverslips were mounted using the Vectashield mounting reagent (<u>Vector Laboratories,</u> VECH-1000, without DAPI).

### Protein aggregate staining using Proteostat^®^

MEFs were grown on coverslips and treated with 5 μg/ml puromycin for 2 h, 100 nM Bafilomycin A1 for 24 h, or subjected to heat stress at 42 °C for 1 h. Cells were fixed in 4% paraformaldehyde (PFA) for 10 min at 37 °C, permeabilized in 0.5% TritonX-100 solution for 5 min, blocked with 5% FBS-PBS at 37°C for 1 h. Proteostat® (Enzo Life Sciences) was prepared according to the manufacturer’s instructions. Cells were stained using Proteostat® at a 1:2000 dilution in the 1x assay buffer provided in the kit for 20 min. The cells were mounted using Vectashield mounting reagent (<u>Vector Laboratories,</u> VECH-1200, containing DAPI) and imaged using confocal microscopy or Airy scan on an LSM900 confocal system.

### Recombinant protein purification of mCherry-p62 and mCherry-p62 S403E S407E

Purification of mCherry-p62 and mCherry-p62 S403E S407E was performed as described previously (Ikeda *et al*., 2023). Briefly, pGEX6P-mCherry-p62 and pGEX6P-mCherry-p62 S403E S407E plasmids were transformed into *E. coli* BL21(DE3) cells (Agilent, 200131). Transformed bacteria were grown in 2xYT media containing 100 μg/ml ampicillin. Protein expression was induced with 0.1 mM IPTG and cultured overnight at 18°C. Cells were harvested and lysed by sonication with SONIFIER 250 (BRANSON) in Buffer A (50 mM HEPES pH 8.0, 500 mM NaCl) supplemented with 1 mM TCEP, 1 mM PMSF, and cOmplete EDTA-free protease inhibitor cocktail (Roche, 11836170001). Cell debris was separated by centrifugation, and the supernatant was applied to COSMOGEL® GST-Accept resin (Nacalai Tesque, 09277-72) in a chromatography column (Bio-Rad, 7321010). The resin was washed with Buffer A, followed by on-column cleavage with GST-tagged PreScission protease for ∼16 h at 4 °C, the cleaved proteins were collected and concentrated to ∼35 μM in Buffer A supplemented with 10% glycerol using VIVASPIN 20, MWCO 30,000 (Sartorius, VS2022). The proteins were flash-frozen in liquid nitrogen and stored at −80°C until use.

### Recombinant protein purification of GST-empty and GST-UBAN×3

Purification of GST-empty and GST-UBAN×3 was performed as described previously (Fennell *et al*., 2020) . Briefly, pGEX4T1 and pGex4T1-ABIN1-UBAN×3 plasmids were transformed into *E. coli* BL21(DE3) cells (Agilent, 200131). Transformed bacteria were grown in LB medium containing 100 μg/ml ampicillin. Protein expression was induced with 0.5 mM IPTG and cultured overnight at 18°C. Cells were harvested, resuspended in GST Buffer (20 mM Tris-HCl, pH 7.5, 10 mM EDTA, pH 8.0, 5 mM EGTA, 150 mM NaCl, 1 mM PMSF, and 0.1% β-Mercaptoethanol), and lysed by sonication using TOMY Ultrasonic Disruptor (UD-100). 0.5% TritonX-100 was added to the lysate, and the cell debris was removed by centrifugation. Equilibrated Glutathione Sepharose™ beads (Cytiva, 17513201) were added to the supernatant and rotated for 1 h at 4°C. The beads were washed with GST buffer containing 0.5% TritonX-100. Bead-immobilized GST-empty or GST- UBAN×3 was washed and resuspended in 20 mM Tris-HCl, pH 7.5 containing 0.1% NaN_3_. The bead-immobilized GST proteins were stored at 4°C until use.

### Recombinant protein purification of LUBAC from insect cells

LUBAC purification was performed as previously described (Rodriguez Carvajal *et al*., 2021). Briefly, Sf9 cells were grown in ESF 921 Insect Cell Culture Medium Protein Free (Expression Systems 96-001-01) at 27°C and infected with baculovirus encoding for human HOIP, His6-human HOIL-1, and Strep (II)-human SHARPIN at a density of 2 × 10^6^ cells/ml. At 72 h post-infection, the cells were harvested, resuspended, and lysed in 100 mM HEPES, 100 mM NaCl, 100 mM KCl, 100 mM Arg, 10 mM MgSO4, and 20 mM imidazole (pH 8.0) supplemented with cOmplete EDTA-free protease inhibitor cocktail (Roche, 11836170001) using the Constant Systems Cell Disruptor at 1.4 kBar. The lysates were then cleared by centrifugation after the addition of 100 mM benzonase and 10 mM PMSF. Cleared lysates were loaded onto a HisTrap FF cartridge (GE Healthcare), proteins were eluted with buffer (100 mM HEPES, 100 mM NaCl, 100 mM KCl, 50 mM arginine, 500 mM imidazole, pH 8.0), and loaded onto a Streptactin Superflow cartridge (IBA Lifesciences 2-1238-001), followed by washes with washing buffer (100 mM HEPES, 100 mM NaCl, 100 mM KCl, pH 8.0), and elution with elution buffer (100 mM HEPES, 100 mM NaCl, 100 mM KCl, 5 mM D-desthiobiotin, pH 8.0). The eluted proteins were concentrated using Centriprep MWCO 50,000 (Merck, 4311). The proteins were flash-frozen in liquid nitrogen and stored at −80°C until use.

### *In vitro* ubiquitination assay

*In vitro* ubiquitination assays were performed as described previously (Rodriguez Carvajal *et al*., 2021). Briefly, reactions were prepared in assay buffer (50 mM HEPES, 150 mM NaCl, 0.5 mM MgSO4, pH 7.5) containing 0.338 μM Ube1 (ubiquitin-activating enzyme 1), 0.338 μM UbcH7 (ubiquitin-conjugating enzyme H7), 0.338 μM LUBAC with HOIL-1 WT or HOIL-1 C460A, 29.0 µM ubiquitin, and with or without 2 mM ATP (Roche, 10519979000)). The reaction was carried out at 37°C for 3 h. For immunoblotting analysis, the reactions were terminated in SDS buffer and boiled at 95°C. For *in vitro* LLPS assays, the reaction mixture was flash-frozen and stored at −80°C until use.

### *In vitro* LLPS assay

mCherry-p62 or mCherry-p62 S403E S407E (20 μM) was mixed with 20% volume of the *in vitro* ubiquitin assay reaction mixture in LLPS buffer containing 20 mM HEPES pH7.5 and 150 mM NaCl (Ikeda *et al*., 2023). The LLPS reaction was performed on a glass-bottom dish coated with 3% BSA. The reaction was incubated at ∼23°C for 1 h before imaging using a Zeiss LSM900 confocal microscope.

### GST pulldown assay

This method has been previously described (Rahighi *et al*, 2009). MEFs were grown in a 10 cm dish to 90% confluence and treated with 5 μg/ml puromycin for 2 h. Following the treatment, cells were lysed in chilled lysis buffer (50 mM HEPES, 150 mM NaCl, 1 mM EDTA, 1 mM EGTA, 25 mM NaF, 10 mM ZnCl2, 10% glycerol, 1% Triton X-100, 20 mM NEM, and Complete protease inhibitor (Roche, 11836170001)). Cell debris was removed at 10,000 rpm, at 4°C for 10 min. 10% of the supernatant was kept aside as input, and the remaining was divided for GST-empty or GST- UBANx3 pulldown. GST-empty or GST-UBAŃ3 coupled beads were incubated with clarified lysates overnight at 4°C. The beads were then washed three times with chilled lysis buffer. The precipitated proteins were analyzed by western blotting.

### Immunofluorescence staining

SH-SY5Y cells transfected with ⍺-Synuclein and pEGFP-C1 human HOIL-1 or pEGFP-C1 human HOIL-1 C460A, as described earlier, were fixed in 4% PFA after 72 h transfection. Cells were permeabilized in 0.5% TritonX-100-PBS for 5 min, blocked with 5% FBS-PBS, and stained with primary antibody for 1 h at 37°C. Samples were washed in PBS and incubated with secondary antibody, Alexa Fluor 568 goat anti-mouse IgG (H+L) (Invitrogen, A11031), or Alexa Fluor 488 goat anti-rabbit IgG (H+L) (Invitrogen, A11008), Alexa Fluor 568 goat anti-rabbit IgG (H+L) (Invitrogen, A11036), Alexa Fluor 488 goat anti-mouse IgG (H+L) (Invitrogen, A11001) Alexa Fluor 488 goat anti-rat IgG (H+L) (Invitrogen, A11006) and Alexa Fluor 405 goat anti-mouse IgG (H+L) (Invitrogen, A31553) for 1 h at 37°C. Nuclei were counterstained with DRAQ7 (Abcam, D15106), as indicated in the figures. Coverslips were mounted using the Vectashield mounting reagent (Vector Laboratories, VECH-1200, containing DAPI, or VECH-1000, without DAPI) and imaged using confocal microscopy or Airyscan on an LSM900 confocal microscope (Zeiss). MEFs were seeded onto cover slips. For staining endogenous proteins, cells were fixed in cold methanol for 10 min at −20 °C, rehydrated in cold PBS on ice, and washed three times. Coverslips were blocked in 5%BSA-PBS incubated with the indicated primary antibodies (anti-LC3 (Nanotools, 0260-100/LC3-2G6), anti-LAMP2 (Abcam, ab13524) anti-p62 (MBL, PM045), anti-α-Synuclein [MJFR1] (Abcam, ab138501), anti-phosphorylated α-Synuclein (Wako, 015-25191), or anti-ubiquitin (FK2) (Enzo, BML-PW8810-0500)) overnight. Samples were stained with the secondary antibody as described above and visualized using an LSM900 confocal microscope (Zeiss).

### 1,6-Hexanediol disruption of condensates in cells

Cells (0.2 ×10^6^) were seeded onto poly-L-lysine (Cosmo Bio, SPL01)-coated coverslips in 6-well plates. The following day, cells were treated with 0%, 5%, and 10% 1,6-Hexanediol (Wako, 087-00432) for 1 min at room temperature after treatment with 5 μg/ml puromycin, as stated in the figure legend (Yu *et al*, 2015). After removing 1,6-Hexanediol, the cells were immediately fixed with 4% PFA for 10 min at 37°C. The cells were washed with PBS, permeabilized in 0.5% TritonX- 100-PBS for 5 min, blocked with 5% FBS-PBS at 37°C for 1 h, and subjected to immunofluorescence as described above.

### Image analysis

Image analysis was performed using FIJI image analysis software (Schindelin *et al*, 2012). To determine the number, size, and properties (e.g., circularity) of the p62 puncta, a threshold was first applied to the images to highlight only the p62-stained particles. Subsequently, the ‘Analyze Particle’ function was applied. To determine α-Synuclein-phospho-α-Synuclein, LC3-p62, or LAMP2-p62 double-positive puncta, a threshold was first applied to the 16-bit images stained with α-Synuclein or p62 to highlight positively stained particles. Regions of interest (ROI) and signal intensities were measured. The same ROI was applied to the 16-bit images stained with the corresponding antibodies (phospho-α-Synuclein, LC3, or LAMP2). The signal intensity in the ROI was measured. Particles with signal intensities greater than the threshold cutoff were defined as double positive. Cells were counted using a threshold value. Watershed was used to separate the overlapping nuclei.

### Sucrose gradient fractionation

SH-SY5Y cells were transfected with ⍺-Synuclein and pcDNA 3.1 human HOIL-1 or pcDNA 3.1 human HOIL-1 C460A. After 72 h of transfection, the cells were lysed in lysis buffer containing 50 mM HEPES, 150 mM NaCl, 1 mM EDTA, 1 mM EGTA, 1% Triton X-100, 10% glycerol, 25 mM NaF, 10 mM ZnCl_2_, 10 mM NEM, 1 mM PMSF, 5 mM Na_3_VO_4_, pH 7.4, supplemented with cOmplete Mini EDTA-free protease inhibitor cocktail (Roche, 11836170001). For sucrose gradient fractionation performed in MEFs, cells were treated with 5 μg/ml puromycin for 2 h, or after recovery for 2 h, control cells were left untreated. Cells were harvested after the indicated treatments and lysed in lysis buffer containing 50 mM HEPES, 150 mM NaCl, 1 mM EDTA, 1 mM EGTA, 1% Triton X-100, 10% glycerol, 25 mM NaF, 10 mM ZnCl_2_, 10 mM NEM, 1 mM PMSF, 5 mM Na_3_VO_4_, pH 7.4 supplemented with cOmplete Mini EDTA-free protease inhibitor cocktail (Roche, 11836170001). The lysates were fractionated in a sucrose gradient from 5% to 25% sucrose through ultracentrifugation at 55,000 rpm, 4°C, for 1.5 h in the Micro Ultracentrifuge (HIMAC Eppendorf Group, CS100FNX).

### Autophagy flux assay using GFP-mCherry-p62

For the autophagy flux assay (Kimura *et al*., 2007; Mizushima & Murphy, 2020), GFP-mCherry-p62 stable MEF lines (*Hoil-1^+/+^*and *Hoil-1^C458A/C458A^*) were grown in a glass-bottom 35 mm dish (Matsunami) to a confluency of 80-90%. The cells were treated with 5 μg/ml puromycin, and 60 frames were imaged with a 2 min interval between frames in the LSM900 confocal system. During imaging, the dishes were kept in a heated hydration chamber with CO_2_ levels maintained at 5%.

### Fluorescence recovery after photobleaching (FRAP)

For Fluorescence recovery after photobleaching (FRAP) analysis, GFP-p62 was stably expressed in *Hoil-1^+/+^*and *Hoil-1^C458A/C458A^* MEFs. The cells were treated with 5 μg/ml puromycin for 2 h before live-cell imaging. Photobleaching was performed with one pulse using a 488 nm laser at 75% intensity within the region of interest (ROI), as indicated in the figure. Live imaging was performed at 5 s intervals for 3 min 20 s. FRAP analysis was performed as described previously (Zheng *et al*, 2011). Fluorescence intensity was measured within the ROI and normalized against the unbleached control region. Mobile fractions (f_m_) and immobile fractions (f_i_) were calculated using the equations f_m_= F_¥_/F_0_, where F_¥_ is the fluorescence at full recovery and F_0_ is the fluorescence before photobleaching, and f_i_= 1- f_m_. The mobile fractions were compared between the HOIL-1 wild type and mutant samples using an unpaired t-test for statistical analysis. The percentage recovery was then plotted.

### Statistics and data reproducibility

Statistical tests between the two groups were performed using a two-tailed unpaired t-test. For non-parametric datasets, the Mann-Whitney test was performed between two sample sets, and the Kruskal-Wallis test was performed for multiple sample sets. Graphical representations and statistical analyses were performed using GraphPad Prism 10.4.2. N for all the experiments are indicated in the figure legends.

## Acknowledgements

We would like to thank all members of the Ikeda Lab for their insightful discussions. We also thank Tomohiro Yamazaki and Tetsuro Hirose (The University of Osaka) for their input on condensate formation assays in cells, and the FBS Core Facility (The University of Osaka) for technical support. This study was supported by JSPS KAKENHI Grant Numbers JP21H04777 (FI), JP23K19350 (SK), JP23K20044, JP24H00060, JP25H01323 (MK), JP24K0197, JP24H01882 (JS), AMED Grant Numbers JP22gm1410004h0003 (MK), JP25gm6910004 (JS), the Takeda Science Foundation (FI), and Project MEET (Osaka University Graduate School of Medicine in association with Mitsubishi Tanabe Pharma Corporation) (FI).

## Disclosure of AI usage

During the preparation of this manuscript, the authors used Paperpal to check grammar. After using this tool, the authors reviewed and edited the content as required and take full responsibility for the content of the published work.

## Author contributions

SK, MM, TN, and HN performed the experiments and analyzed the data; JS and MK provided essential materials and critical advice; and FI conceived and conceptualized the work and strategy. SK and FI drafted and edited the manuscript with inputs from other authors. All authors have read and approved the article.

