## Supplemental information for "Excess Met1-ubiquitination leads to solid aggregate formation"

#### Supplementary figure legends

**Figure S1: Aggregate-prone proteins accumulate in cells expressing HOIL-1 catalytically inactive mutant**

A. Sucrose gradient fractionation in SH-SY5Y cells expressing α-Synuclein aggregates, and HOIL-1 or HOIL-1 C460A. Fractions 1-10, and the lowest fraction (pellet) were loaded, and total cell lysate (TCL) was loaded as a control. The distribution of LUBAC components (HOIP, HOIL-1, and SHARPIN) and α-Synuclein proteins was determined by western blotting analysis. N=2.

B. Co-localization of α-Synuclein aggregates with p-α-Synuclein in *Hoil-1*$^{+/+}$ and *Hoil-1*$^{C458A/C458A}$ MEFs. N=3 independent biological repeats. Scale bar = 10 µm.

C. Graphical representation of the size of α-Synuclein aggregates from (B). Values plotted are mean ± SEM of *n*= >100 cells in each sample set from N=3 independent biological replicates. Mann-Whitney statistical test was performed (****P < 0.0001).

D. Graphical representation of the ratio of p-α-Synuclein/ total α-Synuclein puncta from (B). Values plotted are mean ± SEM of n=12 fields from N=3 independent biological replicates. Unpaired t-test statistical analysis was performed (*P = 0.0268).

**Figure S2: Aggregate accumulation in *Hoil-1*$^{C458A/C458A}$ cells under proteotoxic stresses**

A-C. Representative images of Proteostat-stained aggregates (indicated by arrows) in *Hoil-1*$^{+/+}$ and *Hoil-1*$^{C458A/C458A}$ MEFs under (A) untreated or proteotoxic stress, (B) 100 nM Bafilomycin A1 (BafA1), and (C) heat stress at 42ºC. Scale bar = 5 µm. Graphical representations of the size of the Proteostat-stained aggregates are also shown. Values plotted are mean ± SEM from N=2 independent biological repeats (untreated, *n*= >75 cells; BafA1, *n*= >30 cells; heat stress, *n*= >55 cells per group). Mann-Whitney statistical test was performed; ns, P> 0.05, ****, P< 0.0001, **, P= 0.0029.

**Figure S3: Aggrephagy is compromised in *Hoil-1*$^{C458A/C458A}$ MEFs**

A, B. Fluorescence intensities of p62 (red) and LAMP2 (green) plotted along the yellow arrow to indicate the overlap of p62 and LAMP2 in (A) *Hoil-1*$^{+/+}$ and (B) *Hoil-1*$^{C458A/C458A}$ MEFs. Representative images from N=3 independent biological repeats.

C. Time-lapse imaging to monitor autophagy flux of GFP-mCherry-LC3 *Hoil-1*$^{+/+}$ MEFs and *Hoil-1*$^{C458A/C458A}$ under 5 μg/ml puromycin treatment (*n*= 40 cells from N=3 independent biological repeats). Scale bar = 5 μm.

D. Graphical representation of the ratio of $LC3^{GFP}$/ $LC3^{mCherry}$ fluorescence intensity values in (C). The values plotted are mean ± SD. Mann-Whitney statistical test was performed (****, $P < 0.0001$).

E. qPCR analyses to show that autophagy-related gene expression does not alter significantly between *Hoil-1*$^{+/+}$ and *Hoil-1*$^{C458A/C458A}$ MEFs under 5 μg/ml puromycin treatment. Mean relative fold change ± in SEM from N=3 independent biological replicates were plotted.

**Figure S4: Linear ubiquitin chains may induce alteredproperty of p62 bodies**

A. Graphical representation of all aggregates present in *Hoil-1*$^{C458A/C458A}$ MEFs after recovery, comparing the aggregate circularity with respect to its size. Kruskal-Wallis statistical analysis was performed (****$P < 0.0001$). Data from N>3 independent biological replicates.

B. Representative images demonstrating the dissolution of LLPS condensates of 3×FLAG-FUS R495X protein under 5% and 10% 1,6-Hexanediol (1,6-HD) treatment in HEK293T cells. Scale bar = 10 μm.

C. Immunoblotting analysis of *in vitro* ubiquitination assay comparing ubiquitin chains assembled by LUBAC WT and LUBAC containing HOIL-1 C460A. Met1-linked ubiquitin chains and pan-ubiquitin chains were assessed with anti-Met1 Ub and anti-Ub antibodies, respectively.

**Figure S5 Accumulation of linear ubiquitin chains may be responsible for persistent aggregates.**

A. Representative images of western blotting analysis showing *Otulin* knockdown in sh*Otulin* MEFs compared to sh*Cont* MEFs. Anti-Vinculin antibody was used to monitor loading.

B. GST and GST-UBAN×3 pull down from sh*Cont* and sh*Otulin* MEF lysates followed by immunoblotting analysis. Met1-linked ubiquitin chains (top panel) and pan-ubiquitin (middle panel) levels with and without 5 μg/ml puromycin (2 h) are shown. Ponceau S (bottom panel) was used as the loading control. Anti-OTULIN antibody was used to monitor OTULIN knockdown, and anti-vinculin was used to monitor loading of input samples. Representative data from three independent biological replicates.

C. Comparison of Proteostat-stained aggregates in sh*Cont* and sh*Otulin* MEFs under heat stress at 42 °C. Representative images from N=3 independent biological repeats. Scale bar = 5 μm.

D-E. Graphical representations of aggregate sizes under (D) untreated and (E) heat stress conditions in sh*Cont* and sh*Otulin* MEFs. Values plotted are mean ± SEM from N=3 independent biological replicates. Mann-Whitney statistical test was performed; ns, $P>0.05$, *, $P= 0.0354$.

### Figure S1

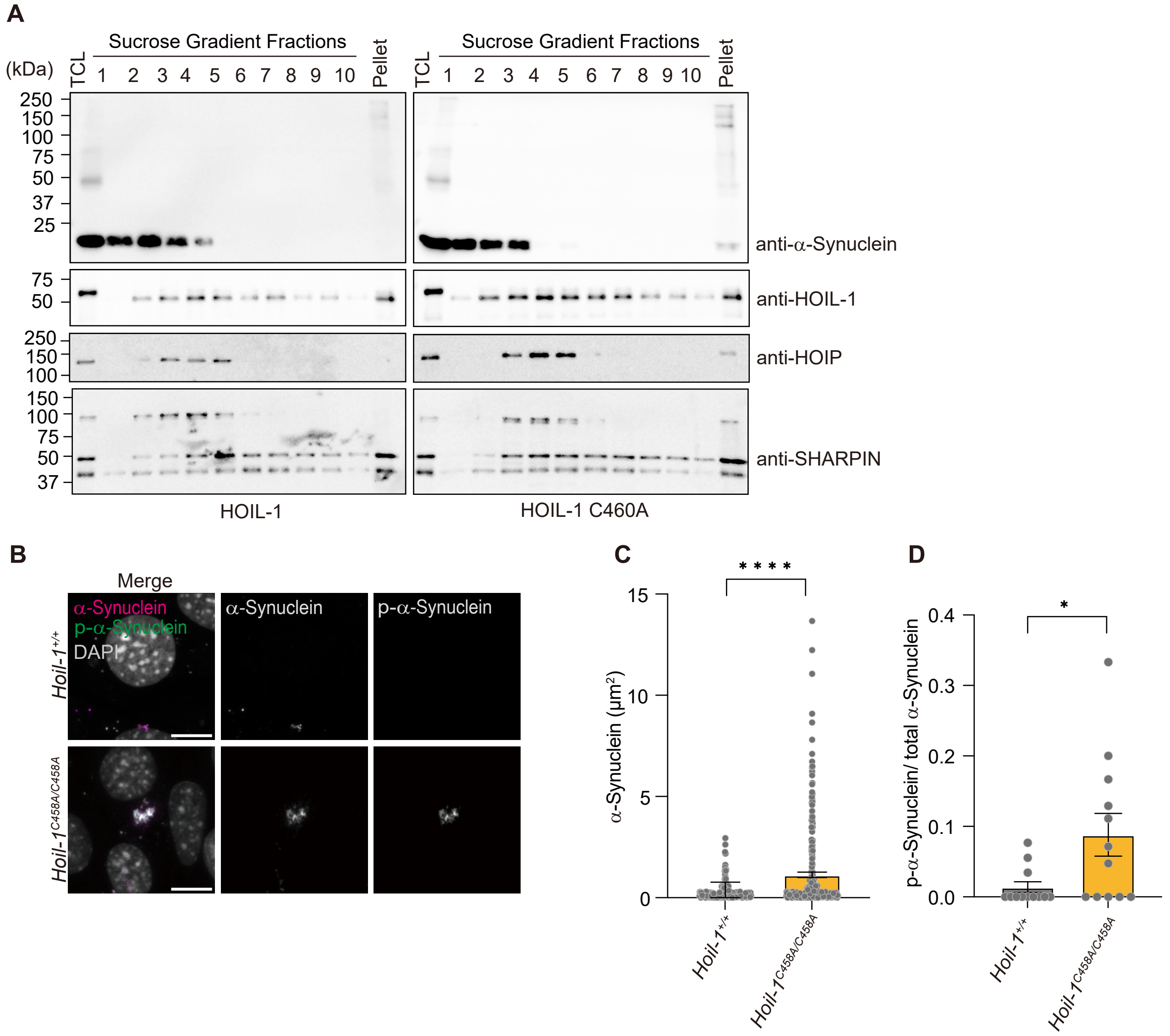

A
Sucrose Gradient Fractions
(kDa)
TCL
1
2
3
4
5
6
7
8
9
10
Pellet
250
150
100
75
50
37
25
anti-α-Synuclein
anti-HOIL-1
anti-HOIP
anti-SHARPIN
HOIL-1
HOIL-1 C460A
B
Merge
α-Synuclein
p-α-Synuclein
DAPI
Hoil-1+/+
Hoil-1C458A/C458A
C
α-Synuclein (μm²)
D
p-α-Synuclein/ total α-Synuclein

### Figure S2

A

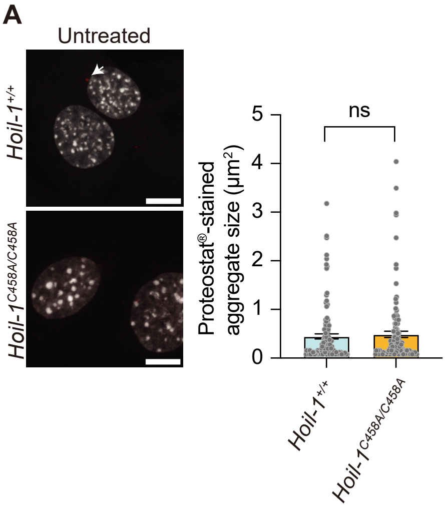


B

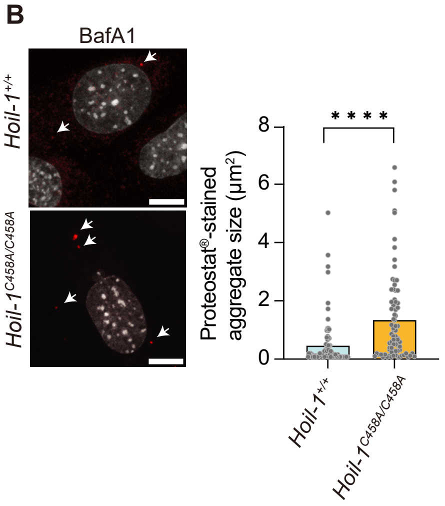


C

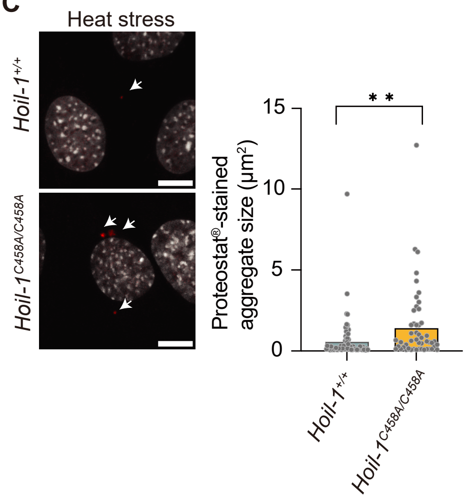

### Figure S3

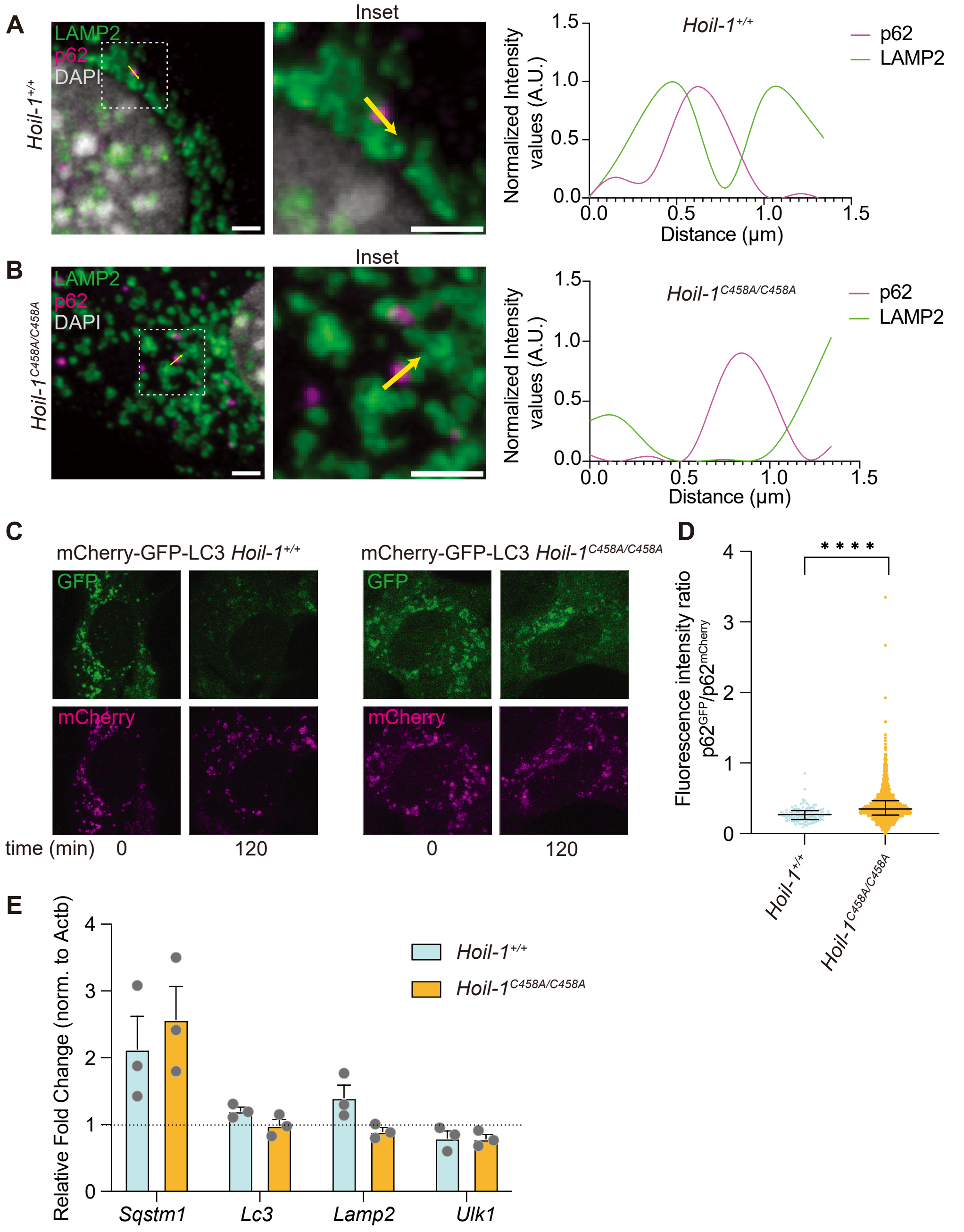

A
LAMP2
p62
DAPI
Hoil-1+/+
Inset
Hoil-1+/+
p62
LAMP2
Normalized Intensity values (A.U.)
Distance (μm)
B
LAMP2
p62
DAPI
Hoil-1C458A/C458A
Inset
Hoil-1C458A/C458A
p62
LAMP2
Normalized Intensity values (A.U.)
Distance (μm)
C
mCherry-GFP-LC3 Hoil-1+/+
mCherry-GFP-LC3 Hoil-1C458A/C458A
GFP
mCherry
time (min) 0 120
0 120
D
Fluorescence intensity ratio p62GFP/p62mCherry
****
Hoil-1+/+
Hoil-1C458A/C458A
E
Relative Fold Change (norm. to Actb)
Hoil-1+/+
Hoil-1C458A/C458A
Sqstm1
Lc3
Lamp2
Ulk1

### Figure S4

A

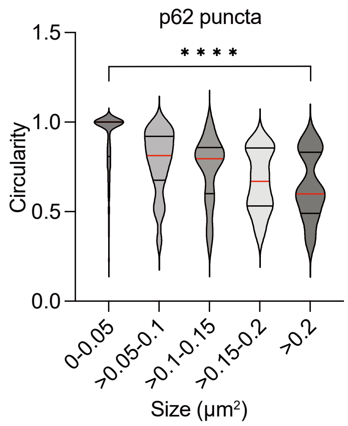


B

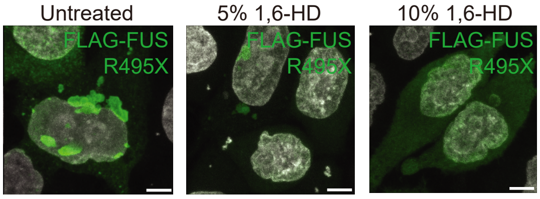


C

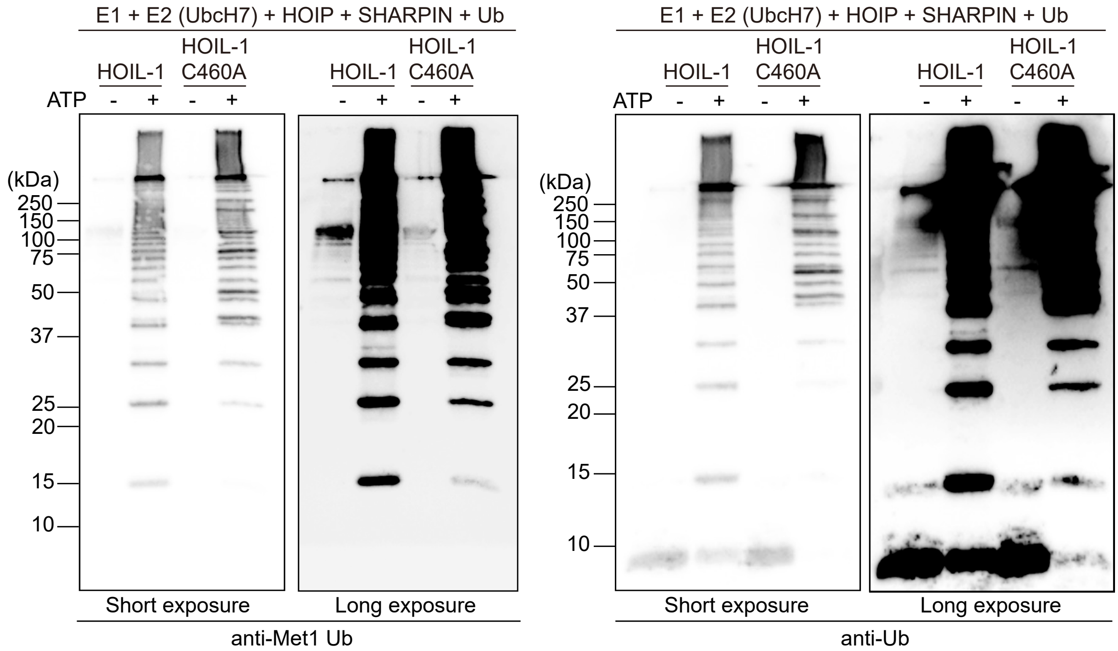

#### Figure S5

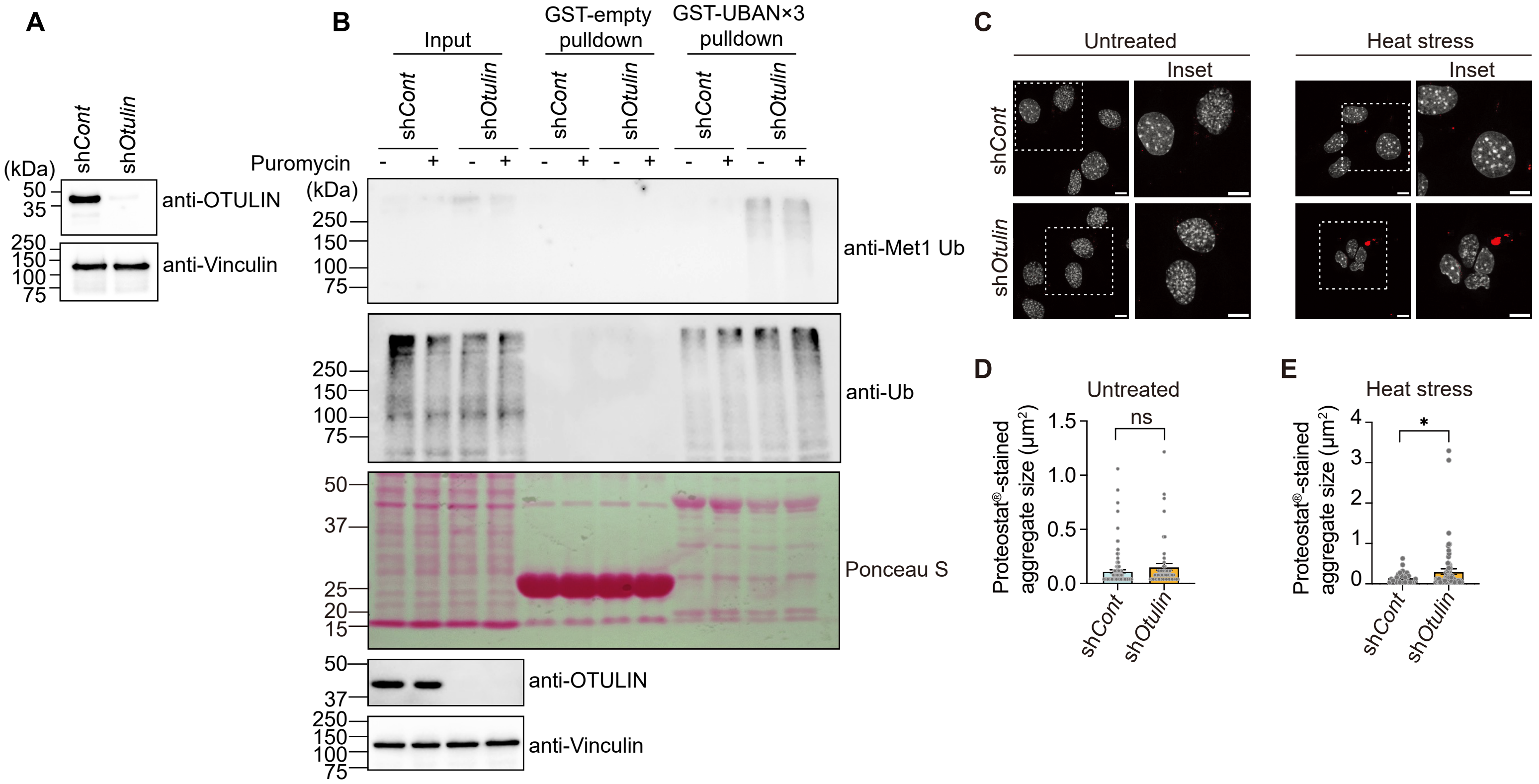
A
(kDa)
shCont
shOtulin
50
35
250
150
100
75
anti-OTULIN
anti-Vinculin
B
Input
GST-empty pulldown
GST-UBAN×3 pulldown
shCont
shOtulin
Puromycin
- +
(kDa)
250
150
100
75
anti-Met1 Ub
anti-Ub
50
37
25
20
15
Ponceau S
anti-OTULIN
anti-Vinculin
C
Untreated
Heat stress
Inset
shCont
shOtulin
D
Untreated
ns
Proteostat®-stained aggregate size (μm²)
1.5
1.0
0.5
0.0
E
Heat stress
*
4
3
2
1
0
